# Increased HCN1 activity in human excitatory neurons drives excessive network bursting in Dravet syndrome

**DOI:** 10.1101/2025.10.03.680281

**Authors:** Federica Riccio, Guilherme Neves, Michelle Gottlieb-Marra, Ariana Gatt, Christina Toomey, Jernej Ule, Ivo Lieberam, Juan Burrone

**Author notes:** Equal contribution (Ivo Lieberam, Juan Burrone).

## Abstract

Dravet syndrome (DS) is a severe childhood epilepsy caused by mutations of the sodium channel NaV1.1. These mutations are thought to compromise the ability of inhibitory interneurons to regulate network activity, leading to seizure events. However, standard treatments to restore inhibition have limited efficacy, suggesting the existence of additional pathological mechanisms. Here, we use hiPSC-derived neuronal networks containing both excitatory and inhibitory neurons to show that excessive bursting activity in DS cultures is driven by excitatory neurons. This rise in bursting frequency is caused by the increased expression of HCN1 “pacemaker” channels in excitatory neurons, and bursting activity can be normalised using a channel blocker. With this work, we propose a new pathophysiological mechanism in DS and identify HCN1 as a novel therapeutic target.

## INTRODUCTION

Dravet syndrome (DS) is a rare form of epileptic encephalopathy characterized by severe pharmacoresistant seizures and comorbidities including developmental delays, motor deficits and behavioural disorders ^1–3^. Most DS cases are caused by loss-of-function mutations in *SCN1A*, the gene encoding for the voltage gated sodium channel NaV1.1 ^4^. How the loss-of-function of a sodium channel, essential for action potential initiation and neuronal activity, paradoxically culminates in hyperexcitable, seizure-prone networks has been a central question in DS pathophysiology.

The “interneuron hypothesis” proposes reduced inhibitory tone (disinhibition) as the mechanistic basis for epileptic activity in DS. This theory, based on the preferential expression of *SCN1A* by GABAergic inhibitory interneurons ^5^, is supported by more than a decade of research in DS mouse models, showing that reduced Na^+^ currents and hypoexcitability of interneurons lead to network hyperexcitation ^6–9^. However, recent studies in mouse models ^10,11^ and, particularly, human induced pluripotent stem cell (hiPSC)-derived neurons ^12–14^, are starting to challenge the interneuron hypothesis and propose increased excitatory activity as a parallel pathological mechanism in DS. Crucially, the relative contribution of excitatory and inhibitory dysfunction to network hyperactivity in DS remains largely unexplored in human neurons. Furthermore, the molecular mechanisms driving a gain-of-function hyperactivity phenotype in excitatory neurons with Na^+^ channel loss-of-function mutations, remains unknown.

In this study, we leveraged the use of hiPSC-derived excitatory/inhibitory co-cultures to dissect the contribution of each neuronal type to DS pathophysiology. We showed that, although both neuronal types contribute to network hyperactivity, it is the excitatory neurons that play a dominant role. DS excitatory neurons showed increased expression of the “pacemaker” channel HCN1, which is responsible for regulating rhythmic firing patterns and intrinsic bursting in neurons as well as cardiac muscle cells ^15,16^. We identified the increase in the current conducted by HCN1 channels (*I*_h_) as the molecular mechanism responsible for the excessive bursting activity displayed by DS excitatory neurons and showed that blocking HCN1 activity is sufficient to reduce network hyperactivity to control levels. Our findings have uncovered the main neuronal type and molecular mechanism responsible for epileptic activity in Dravet syndrome human networks, with important implications for the development of novel therapeutic approaches.

## RESULTS

### Increased neuronal activity in Dravet syndrome patient-derived cortical networks

To investigate DS phenotypes in the context of human cortical networks, we used hiPSC-derived excitatory and inhibitory neurons to generate a ratio-defined, cortical-like, co-culture system (Figure. 1A). Three hiPSC lines derived from DS patients (DS1, DS2 ^12^ and DS5 ^17^) and two control lines (Wildtype and IsoDS1) were used. IsoDS1 is an isogenic control line generated in this study by correcting the *SCN1A* mutation in the DS1 line using CRISPR technology (Figure S1A-D). Each hiPSC line was engineered to carry the doxycycline inducible transcription factors necessary for forward programming into excitatory or inhibitory neurons and express the genetically encoded calcium indicators GCaMP6f or jrGECO1a, respectively (Figure 1A and Figure S1E-F). Following well established protocols ^18,19^, we derived excitatory and inhibitory neurons that express multiple cortical markers as well as neuronal subtype-specific genes, fire repetitive action potentials, produce voltage dependent currents and form excitatory and inhibitory synapses (Figure S2A-D).

**Figure 1.**
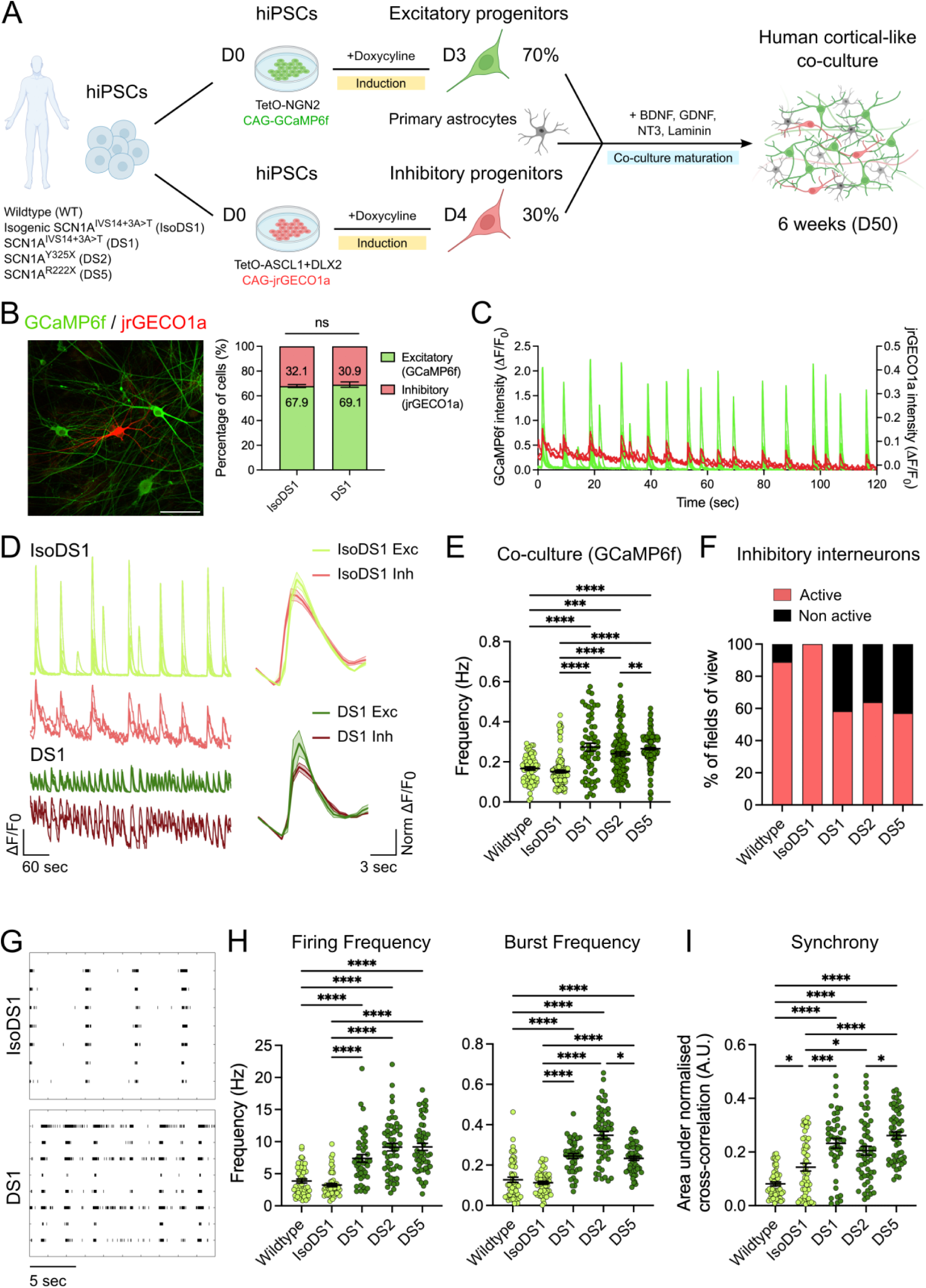
Dravet syndrome patient-derived excitatory/inhibitory cortical networks display a pathological increase in neuronal activity. (**A**) A schematic of the hiPSC lines used in this study, their forward programming differentiation into excitatory and inhibitory progenitors and their assembly into a mature, cortical-like, co-culture. (**B**) Left, fluorescence image of a co-culture of GCaMP6f-expressing excitatory neurons and jrGECO1a-expressing inhibitory interneurons stained for GFP (green) and RFP (red). Scale bar 100 µm. Right, quantification of the cell type ratio at 6 weeks of maturation (D50) plotted as the percentage of excitatory (green) and inhibitory (red) cells in IsoDS1 and DS1 isogenic co-cultures. Mean ± SEM, Two-way ANOVA with multiple comparison, ns = not significant (p>0.05), n=3 independent experiments (12 fields of view). (**C**) Representative calcium traces from excitatory (GCaMP6f – green) and inhibitory (jrGECO1a – red) neurons in a IsoDS1 co-culture at 6 weeks (D50), plotted as the ΔF/F_0_ over time. (**D**) Left, representative calcium traces from excitatory (green) and inhibitory (red) neurons in IsoDS1 and DS1 co-cultures at 6 weeks (D50), plotted as the ΔF/F_0_ over time (60 sec). Right, event-triggered average calcium traces for excitatory (green) and inhibitory (red) neurons in IsoDS1 and DS1co-cultures. Mean ± SEM. The trigger event is the network burst and the amplitude for each cell type is normalised to the average amplitude of all calcium events detected for the same cell. (**E**) Quantification of calcium signal frequency recorded for excitatory neurons (GCaMP6f) in co-culture at 6 weeks (D50). Mean ± SEM, Kruskal-Wallis test with Dunn’s multiple comparison, **p < 0.01, ***p < 0.001, ****p < 0.0001, n=3/4 independent experiments (58-176 cells). (**F**) Quantification of the percentage of analysed fields of view containing active inhibitory interneurons in co-culture at 6 weeks (D50). (**G**) Representative raster plots showing 20 sec of activity exhibited by IsoDS1 and DS1 co-culture networks at 6 weeks (D50). (**H-I**) Quantification of the firing frequency, burst frequency and synchrony recorded during week 6 of co-culture. Mean ± SEM, Kruskal-Wallis test with Dunn’s multiple comparison, *p < 0.05, ***p < 0.001, ****p < 0.0001, n=3 independent experiments (each with 6 wells per genotype, 3 recordings – D42, D46, D50).

To generate co-culture networks, excitatory and inhibitory progenitors were mixed at a ratio of 70:30 to mimic the ratio found in the human cortex ^20,21^ (Figure 1A-B). After 6 weeks of maturation, we used two-colour calcium imaging to simultaneously assess both excitatory and inhibitory activity within the networks. We observed synchronous firing of excitatory and inhibitory neurons in co-culture, both in control and DS conditions, suggesting the formation of a functionally integrated network (Figure 1C-D). The frequency of calcium events was significantly increased in all three DS co-cultures (Figure 1E), however, we found that only ∼60% of the fields of view analysed for the DS co-cultures contained active inhibitory neurons, compared to 90-100% for the controls (Figure 1F). These observations indicate a hyperactive nature of DS networks associated with impaired activity of inhibitory interneurons.

Surprisingly, we also observed an unexpected decrease in the amplitude of calcium events in excitatory neurons of DS cultures (Figure S3A). To explore this further, we carried out simultaneous electrophysiological recordings and calcium imaging in the same neuron. We found that each calcium event was driven by a burst of action potentials (APs) (Figure S3B-C), with DS1 neurons showing a similar number of spikes per burst (Figure S3D) but a higher spike frequency within burst (lower inter-spike-interval; Figure S3E), when compared to IsoDS1 neurons. Characterization of calcium signals in response to field stimulation at different frequencies showed an inverse relation between calcium signal amplitude and stimulation frequency that was independent of genotype (Figure S3F). Similar findings have been previously reported in modelling studies ^22,23^. We conclude that the decrease in the amplitude of calcium events in DS1 neurons is likely a consequence of higher spike frequencies during a burst. Since the calcium signal amplitude reflects a non-linear aggregation of number of spikes per burst, burst duration and spike frequency within burst, we focused exclusively on calcium signal frequency for the rest of this study.

To further explore differences in network activity with higher temporal resolution, we grew co-culture networks on Multi Electrode Arrays (MEAs). Consistent with our calcium imaging data, we observed a significant increase in mean firing frequency, burst frequency and network burst frequency in all three DS co-cultures after 6 weeks (Figure 1G-H and Figure S3G). We also found an increase in the frequency of spikes within a burst, but not in the number of spikes per burst (Figure S3H-I), which is in line with the observations we made when performing simultaneous calcium imaging and patch-clamp recordings (Figure S3D-F). Finally, MEA recordings showed a higher degree of synchrony in the DS co-cultures compared to controls (Figure 1I). Overall, all three Dravet co-cultures displayed different aspects of neuronal and network activity that are consistent with an epileptic-like phenotype. The most significant feature of this phenotype is the increase in burst frequency, expressed at both the single neuron and network level (Figure S3J). Notably, bursts are also shorter but carry spikes at a higher frequency compared to controls (Figure S3J).

### Excitatory neurons contribute to increased network activity in Dravet syndrome

The epileptic-like activity observed in Dravet networks could be the result of impaired inhibition, excessive excitation, or a combination of the two. To assess the relative contribution of excitation and inhibition towards the overall phenotype observed in DS co-cultures, we generated mixed genotype co-cultures. For each disease line, we generated co-cultures of either wildtype excitatory neurons grown together with Dravet inhibitory interneurons, or Dravet excitatory neurons grown together with wildtype inhibitory interneurons (Figure 2A), and compared them to single genotype co-cultures.

**Figure 2.**
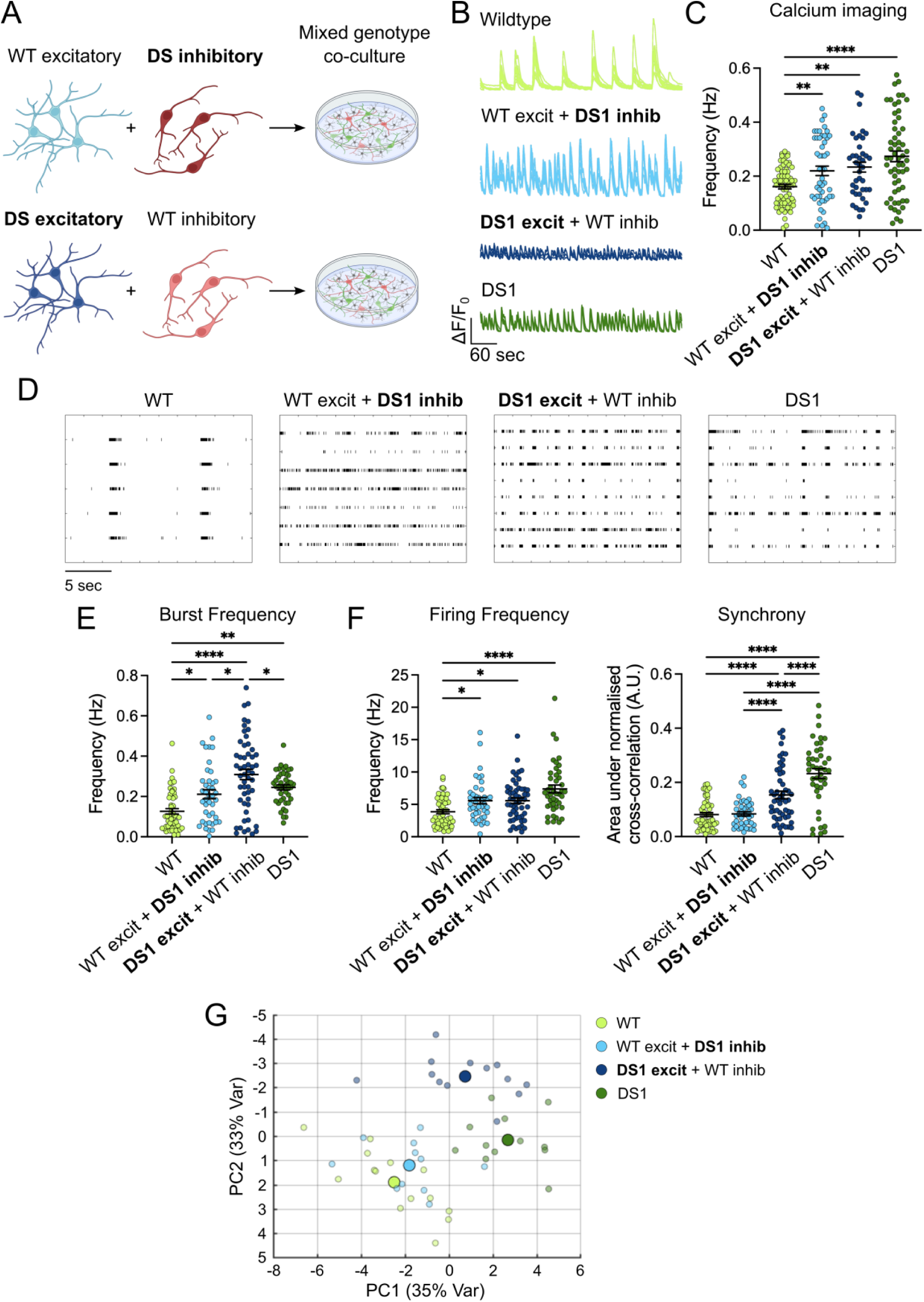
Excitatory neurons are the main neuronal type contributing to pathological hyperactivity of patient-derived Dravet networks. (**A**) A schematic of the mixed genotype co-cultures generated for this study. (**B**) Representative calcium traces from excitatory neurons at D50 in different co-culture conditions: WT excitatory + WT inhibitory, WT excitatory + DS1 inhibitory, DS1 excitatory + WT inhibitory, DS1 excitatory + DS1 inhibitory. Calcium signal is plotted as the ΔF/F_0_ over time (60 sec). (**C**) Quantification of calcium signal frequency recorded for excitatory neurons (GCaMP6f) in mixed– and same-genotype co-cultures at 6 weeks (D50). Mean ± SEM, Two-way ANOVA with Tukey’s multiple comparison, **p < 0.01, ****p < 0.0001, n=3 independent experiments (40-69 cells). (**D**) Representative raster plots showing 20 sec of activity for each co-culture conditions at 6 weeks (D50). (**E-F**) Quantification of the burst frequency, firing frequency and synchrony recorded in the different co-culture conditions during week 6. Mean ± SEM, Two-way ANOVA with Tukey’s multiple comparison, *p < 0.05, **p < 0.01, ****p < 0.0001, n=3 independent experiments (each with 6 wells per genotype, 3 recordings – D42, D46, D50). (**G**) Principal Component Analysis (PCA) of electrophysiological parameters, listed in the methods section, extrapolated from field potential recordings (MEAs) of mixed– and same-genotype co-cultures at week 6 (D42, D46, D50). Smaller dots represent individual wells, bigger dots represent the average per condition.

An increase in the frequency of calcium events, representing AP bursts, was observed in mixed co-cultures containing DS1 and DS2 excitatory neurons (Figure 2B-C and Figure S4A). In contrast, only mixed co-cultures containing DS1 inhibitory interneurons, but not DS2 or DS5 inhibitory interneurons, showed increased calcium signal frequency (Figure 2B-C and Figure S4A). These results suggest that DS excitatory neuron activity provides a stronger contribution, compared to their inhibitory counterpart, to determine the increase in network activity observed in DS co-cultures. Similar results were observed with MEAs (Figure 2D). Bursting frequency was always increased in mixed co-cultures containing excitatory neurons from DS genotypes, but was mostly unaffected in mixed co-cultures containing inhibitory interneurons from DS genotypes (Figure 2E and Figure S4B). In contrast, mean firing frequency and network synchrony were affected either by the presence of DS excitatory neurons or DS inhibitory interneurons or both, depending on the DS genotype (Figure 2F and Figure S4C-D). Together, these findings show that excitatory and inhibitory neurons both contribute, but in different ways, to the overall epileptic-like phenotype observed in DS networks. While both DS excitatory and inhibitory neurons can affect the mean firing frequency and synchrony of the network, DS excitatory neurons are solely responsible for the increased burst frequency of DS networks. Contrary to the interneuron hypothesis, these observations suggest that the dysregulation of excitatory neurons is a key driving force behind the detected epileptic-like phenotype. In fact, Principal Component Analysis (PCA) of all the MEA parameters showed that, whereas mixed co-cultures containing DS inhibitory interneurons clustered closely with fully wildtype co-cultures, mixed co-cultures containing DS excitatory neurons did not (Figure 2G and Figure S4E), suggesting they provide a larger phenotypic contribution compared to their inhibitory counterpart. Of note, fully DS co-cultures form a third distinct cluster, indicating a synergistic effect of both impaired excitation and inhibition in the disease model.

### Monocultures of Dravet syndrome excitatory neurons show increased bursting activity

Having identified excitatory neurons as the main contributors to the epileptic-like phenotype of DS networks in co-culture, we reasoned that monocultures of DS excitatory neurons, grown without inhibitory interneurons, would show a similar hyperactivity and provide evidence for the cell-autonomous nature of this phenotype (Figure 3A). This would confirm our findings in co-cultures and also facilitate studies aimed at uncovering the underlying mechanisms of excitatory neuron hyperactivity. Both calcium imaging and MEA recordings of excitatory neuron monocultures showed a clear increase in bursting frequency in DS1 and DS2 neurons, but less so in DS5 neurons (Figure 3B-F). Although other MEA parameters also changed, such as the number and frequency of spikes per burst and the level of synchrony within the network (Figure S5A-D), we focused on burst frequency, which is the main readout of network hyperactivity that we observed in the co-cultures. To probe the mechanism of this circuit hyperactivity, we first carried out whole-cell patch-clamp recordings to assess the intrinsic excitability of excitatory neurons. Voltage-clamp recordings revealed only a small increase in the amplitude of a fast inward current (predominantly representing Na^+^ currents), which was only significant for DS1 and DS5 neurons compared to wildtype (Figure S5E). A similar small increase in outward (K^+^) currents was also observed (Figure S5F). These minor changes in voltage-gated conductance did not alter the intrinsic excitability of neurons measured in current-clamp recordings (Figure S5G), nor the shape or properties of APs (Figure S5H-I). Nevertheless, measures of spontaneous firing clearly showed an increase in the frequency of spontaneous bursts (Figure 3G-H and Figure S5J), but not in overall firing rate (Figure S3K), suggesting a shift in the firing patterns of DS excitatory neurons to increased bursting activity, in line with our calcium and MEA recordings. In general, the phenotypes observed in excitatory monocultures were stronger in the DS1 and DS2 lines, compared to the DS5 line. This may in part reflect the well described symptomatic heterogeneity reported in patients with DS ^24–28^.

**Figure 3.**
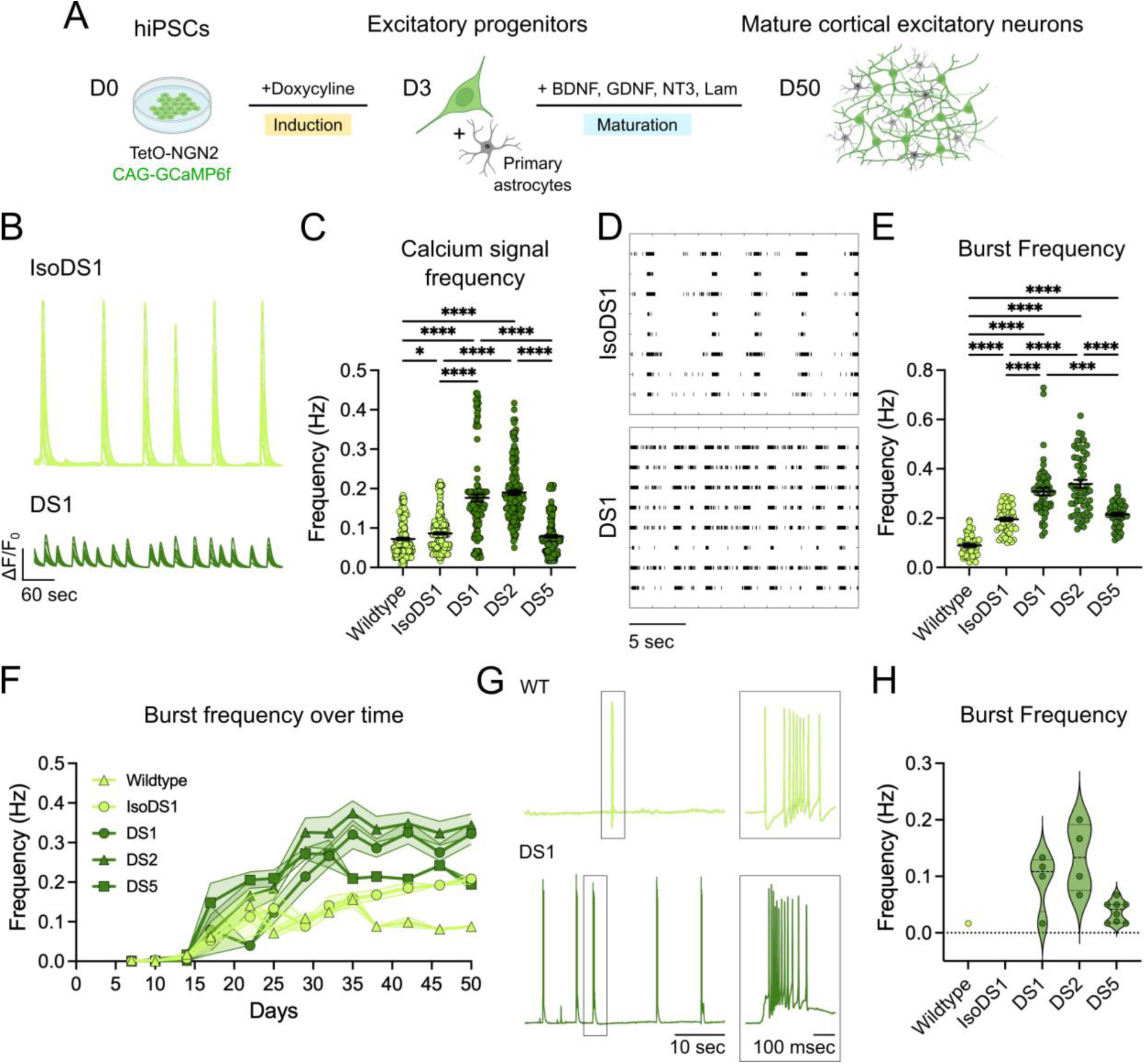
Monocultures of Dravet patient-derived excitatory neurons show increased bursting activity. (**A**) A schematic of the forward programming differentiation into mature cortical excitatory neurons. (**B**) Representative calcium traces from IsoDS1 and DS1 excitatory neuron monocultures at 6 weeks (D50), plotted as the ΔF/F_0_ over time (60 sec). (**C**) Quantification of calcium signal frequency recorded for excitatory neuron monocultures at 6 weeks (D50). Mean ± SEM, Kruskal-Wallis test with Dunn’s multiple comparison, *p < 0.05, ****p < 0.0001, n=3 independent experiments (115-215 cells). (**D**) Representative raster plots showing 20 sec of activity for IsoDS1 and DS1 excitatory neuron monocultures at 6 weeks (D50). (**E**) Quantification of the burst frequency recorded in excitatory neuron monocultures during week 6. Mean ± SEM, Kruskal-Wallis test with Dunn’s multiple comparison, ***p < 0.001, ****p < 0.0001, n=3 independent experiments (each with 6 wells per genotype, 3 recordings – D42, D46, D50). (**F**) Burst frequency over time recorded for excitatory neuron monocultures. Mean ± SEM, n=3 independent experiments (each with 6 wells per genotype). (**G**) Representative current-clamp traces of spontaneous bursting activity recorded for wildtype and DS1 excitatory neurons. Box, detailed view of a burst for each trace. (**H**) Quantification of the burst frequency recorded by current-clamp for *actively bursting* excitatory neuron (fig. S5J) at 6 weeks.

As expected, monocultures of control excitatory neurons grown on MEAs showed higher firing frequencies when compared to control co-cultures that included inhibitory interneurons. However, this difference was either diminished or absent in DS co-cultures, where firing frequency remained equally high despite the presence of interneurons (Figure S5L), providing evidence for some level of interneuron dysfunction in DS networks. In line with this, calcium imaging in monocultures of only inhibitory interneurons showed fewer bursting events in DS lines (Figure S6A-D). Surprisingly, and in contrast with previous reports in the literature ^29,30^, electrophysiological recordings did not reveal a deficit in the amplitude of Na^+^ currents, or other currents, in DS inhibitory interneurons (Figure S6E). Although intrinsic excitability was largely unaffected in DS1 and DS2 interneurons, DS5 interneurons did show a decrease in excitability, particularly to large current injections, and impaired AP properties (Figure S6F-I). Together, our findings confirm a role for both excitatory and inhibitory neurons in Dravet syndrome hyperactivity, but the mechanism does not appear to involve clear-cut changes in intrinsic excitability, particularly for excitatory neurons.

### Increased HCN1 expression and HCN1-mediated currents in Dravet syndrome patient-derived excitatory neurons

Our findings show that excitatory neurons drive excessive bursting frequency in Dravet networks but the mechanisms that lead to this paradoxical “gain-of-function phenotype” following a NaV1.1 loss-of-function mutation is not easily explained. To uncover the molecular mechanism underlying this phenotype, we investigated differential gene expression of excitatory neurons by bulk RNAseq. Although PCA analysis showed that Dravet samples clearly clustered separately from controls (Figure S7A), DS5 samples clustered away from DS1 and DS2, possibly because of the different sex of the DS5 donor (female; Figure S7B). Crucially, the clustering was not influenced by batch preparation (Figure S7C). Gene expression analysis revealed the differential expression of numerous genes, including multiple epilepsy-associated genes (Figure 4A). Amongst these genes was Hyperpolarization Activated Cyclic Nucleotide Gated Potassium Channel 1 (*HCN1*), which is responsible for the generation of the hyperpolarization-activated current (h current; *I*_h_). This channel regulates membrane potential and thereby spontaneous rhythmic activity and bursting in the brain, particularly in cortical and hippocampal neurons, as well as the heart, in cardiac muscle cells of the sinoatrial and atrioventricular node ^15,16^. HCN1 transcript levels were increased in all DS excitatory neurons (Figure 4B). Notably, we observed that the start of HCN1 overexpression coincided with the onset of SCN1A expression (around day 20) and preceded the emergence of the bursting phenotype by only a few days (Figure S7D), suggesting a causal role of HCN1 overexpression in driving the increase in bursting activity. Although overall HCN1 protein levels were also increased in all DS excitatory neurons (Figure 4C-D and Figure S7E), we found that DS1 and DS2 neurons, but not DS5 neurons, showed HCN1 enrichment in dendrites compared to the soma (Figure 4D-F and Figure S7E), a feature that has been associated with neuronal hyperexcitability. Crucially, a similar dendritic enrichment was observed in postmortem brain tissue from DS patients. We saw robust labelling of HCN1 in the hippocampus and cortex in both control and DS patient tissue (ranging from 1 to 6 years old). However, dendritic staining was more apparent in Dravet cases, particularly in the hippocampus where somatic and dendritic compartments are nicely stratified (Figure 4F and Figure S7F). Although quantification of HCN1 subcellular distribution is noisy in postmortem samples, we found that segmentation of dendrites stained for HCN1 was more prominent in Dravet brains (Figure 4F).

**Figure 4.**
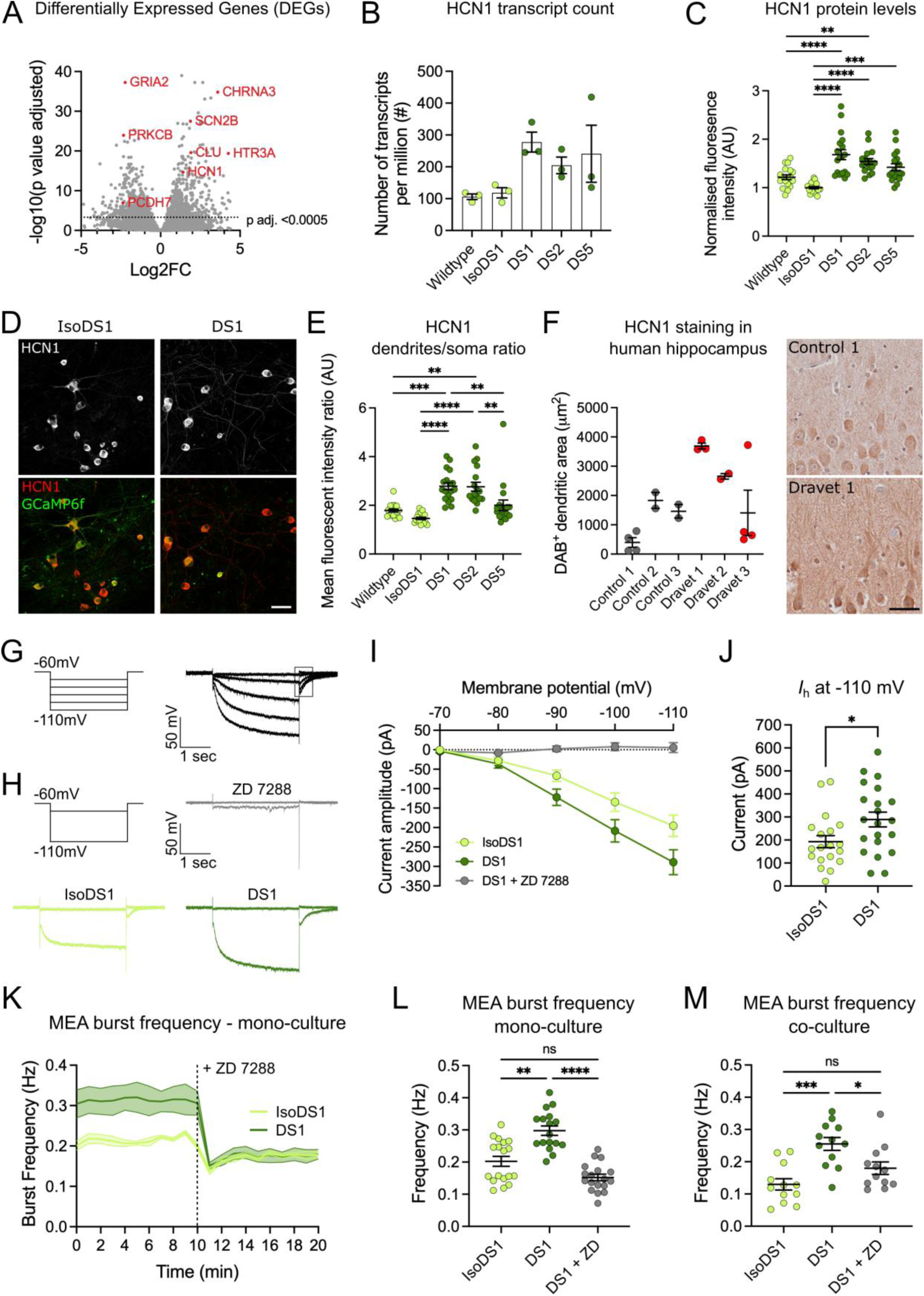
Increased HCN1 expression in Dravet patient-derived excitatory neurons drives the pathological increase in bursting frequency. (**A**) Volcano plot showing differential gene expression in Dravet (DS1, DS2 and DS5) vs. Control (wildtype and IsoDS1) excitatory neurons at 6 weeks (D50). Highlighted in red are some differentially expressed genes with known association to epilepsy. (**B**) HCN1 transcript count from bulk RNA-seq in excitatory neurons at 6 weeks (D50). Mean ± SEM, n=3 independent experiments. (**C**) Quantification of total HCN1 protein levels, measures as HCN1 mean fluorescence intensity within TUBB3^+^ area, normalised for the control condition IsoDS1. Mean ± SEM, One-way ANOVA with Tukey’s multiple comparison, **p < 0.01, ***p < 0.001, ****p < 0.0001, n=3 independent experiments (17-18 fields of view). (**D**) Fluorescence images of IsoDS1 and DS1 excitatory neurons at 6 weeks (D50) stained for HCN1 (red), GCaMP6f (green) and TUBB3. Scale bar 50 µm. (**E**) Quantification of HCN1 dendrites/soma content ratio. Mean ± SEM, Kruskal-Wallis test with Dunn’s multiple comparison, **p < 0.01, ***p < 0.001, ****p < 0.0001, n=3 independent experiments (17-18 fields of view). (**F**) Right, quantification of HCN1^+^ dendritic area in human hippocampal slices from 3 control and 3 Dravet syndrome cases. Mean ± SEM, Welch’s t-test, *p < 0.05 (2-4 slices per case). Left, representative images of human hippocampal slices from one control and one Dravet syndrome case stained for HCN1. Scale bar 50 µm. (**G**) Left, voltage protocol for the recording of h-currents (*I*_h_). Right, representative h-current trace with tail current highlighted by a black box. (**H**) Top left, voltage protocol for *I*_h_ recording showing the first and last voltage steps. Top right, first and last voltage steps recorded from a DS1 excitatory neuron treated with 25µM ZD 7288. Bottom left, first and last voltage steps recorded from a IsoDS1 excitatory neuron. Bottom right, first and last voltage steps recorded from a DS1 excitatory neuron. All recordings were acquired at 6 weeks. (**I**) Quantification of the average *I*_h_ amplitude at each voltage step recorded for IsoDS1 and DS1 excitatory neurons as well as DS1 excitatory neurons treated with 25µM ZD 7288. Mean ± SEM (6-21 cells). (**J**) Quantification of *I*_h_ amplitude at the last voltage step (−110mV) for IsoDS1 and DS1 excitatory neurons at 6 weeks. Mean ± SEM, Welch’s t-test, *p < 0.05 (19-21 cells). (**K**) Representation of burst frequency over time (20 min) for IsoDS1 and DS1 excitatory neurons with addition of 25µM ZD 7288 at 10 min. Mean ± SEM (6 wells). (**L**) Quantification of the burst frequency recorded for IsoDS1 and DS1 excitatory neuron as well as DS1 excitatory neurons treated with 25µM ZD 7288 at D50. Mean ± SEM, Kruskal-Wallis test with Dunn’s multiple comparison, ns = not significant (p>0.05), **p < 0.01, ****p < 0.0001, n=3 independent experiments (each with 6 wells per genotype). (**M**) Quantification of the burst frequency recorded for IsoDS1 and DS1 co-cultures as well as DS1 co-culture treated with 25µM ZD 7288 at D50. Mean ± SEM, One-way ANOVA with Tukey’s multiple comparison, ns = not significant, *p < 0.05, ***p < 0.01, n=3 independent experiments (each with 6 wells per genotype).

To provide a functional readout of HCN1 overexpression, we carried out voltage-clamp recordings to measure *I*_h_ in DS1 and IsoDS1 excitatory neurons (Figure 4G). We found that, on average, *I*_h_ amplitude was increased in DS1 neurons compared to IsoDS1 (Figure 4G-J), as was the *I*_h_ tail current (Figure S8A). Crucially, the currents we measured could be completely blocked with the application of the HCN channel inhibitor ZD-7288 (ZD) ^31–34^ (Figure 4G-I and Figure S8B). Although ZD is not specific for HCN1 over the other HCN channels (HCN2/3/4), our RNAseq dataset showed that only HCN1, but not HCN2/3/4, is expressed in our excitatory neurons (Figure S8C). To test the effects of HCN1 overexpression on neuronal activity, we treated excitatory neurons grown on MEAs with ZD during field potential recordings. Following ZD application, the burst frequency of DS1 excitatory neurons decreased to the same level of IsoDS1 neurons (Figure 4K-L). A small drop in burst frequency was also observed for IsoDS1 neurons, as expected for control neurons that also express HCN1 channels but at a lower level. A significant decrease in bursting frequency was also observed for DS2 excitatory neurons, following ZD treatment, but not for DS5 neurons (Figure S8D-E). This is not surprising considering that the bursting frequency of DS5 excitatory neurons is not significantly increased compared to controls at this time point (Figure 3). Importantly, the normalization of burst frequency was also observed when ZD was applied to excitatory and inhibitory co-cultures (Figure 4M). Although ZD was reported to inhibit some sodium channels ^35^, we found that, at the concentrations used in our experiments, sodium currents were unaffected (Figure S8F), providing further evidence that the reduction in burst frequency by ZD was through direct inhibition of HCN channels. Together, these experiments provide evidence of pharmacological phenotypic rescue of aberrant burst frequency in DS excitatory neurons by HCN1 inhibition.

## DISCUSSION

Our findings challenge the current notion that Dravet syndrome is a disease driven exclusively by interneuron hypoexcitability by showing that patient-derived excitatory neurons not only play a central role in the pathology but that targeting excitatory dysfunction is sufficient to rescue disease phenotypes *in vitro*. While most prior work has focused on intrinsic excitability deficits in inhibitory interneurons as the cause of DS pathology ^6–9,29,30,36^, in this study we show that excitatory neurons, even in the absence of inhibitory impairment, can autonomously develop a pathological hyperactive phenotype (Figure 2 and 3). We show that this hyperactivity is not explained by gross changes in intrinsic excitability but rather by altered burst dynamics driven by the overexpression of HCN1 (Figure 4). These results complement previous studies that have suggested either normal or modestly altered intrinsic excitability in excitatory neurons ^5–7,9–14,37–40^, but have overlooked changes in firing patterns as a pathological feature.

Although the link between the loss-of-function of SCN1A and the increased expression of HCN1 remains unknown, it is possible that initial subtle changes in neuronal activity caused by the loss-of-function of SCN1A could be responsible for driving changes in HCN1 expression. Indeed, the expression of HCN1 is known to be broadly modulated by neuronal activity ^41–45^. In particular, the experimental induction of a single febrile seizure has been shown to cause long-term upregulation of *I*_h_ in pyramidal cell dendrites ^46^, resembling the dendritic enrichment of HCN1 reported here. While enhanced *I*_h_ is typically thought to be associated with decreased excitability ^47–50^, both experimental and computational models have showed that increases in *I*_h_ result in a general enhancement of dendritic excitability ^45,46^. Indeed, recent studies show that gain-of-function mutations in the HCN1 gene itself cause “Dravet-like epilepsy syndromes” ^51–53^ with similar pharmacological sensitivities. Intriguingly, drugs that are broadly characterised as sodium channel blockers (e.g. lamotrigine) and which have been shown to worsen symptoms in both SCN1A and HCN1 related epilepsies ^54,55^, have also been proposed to act as agonists for HCN channels ^56–60^, providing an alternative explanation for their paradoxical effect.

A primary role for HCN1 in the pathophysiology of DS could also explain the reduced severity of the epileptic phenotypes displayed by DS mouse models ^61^, where there is currently little evidence for an involvement of excitatory neurons. These inconsistencies between human patients and mouse models may be explained by the fact that the expression patterns of HCN1 in the brain differ substantially between species. While rodents display low HCN1 expression in superficial cortical layers ^62–64^ and only limited signs of *I*_h_ across various cortical regions ^65–67^, HCN1 is ubiquitously expressed in human supragranular pyramidal neurons and *I*_h_ substantially contributes to the physiology of these neurons ^68^. As a result, aberrant HCN1 activity may only be significant in human excitatory neurons and this could explain why excitatory contributions to DS pathology are more apparent in human iPSC-derived neurons than in rodent models. Nonetheless, recent work in a DS mouse model measured an increase in sag potential in CA1 hippocampal neurons, a feature that may indicate some contribution of pyramidal cells, and HCN1 upregulation, in a DS mouse model ^69^.

Our results argue for a different therapeutic approach in DS, shifting the focus that is currently exclusively on interneurons to favour strategies that also target excitatory neurons. Importantly, we demonstrate that pharmacological inhibition of HCN1-mediated currents normalises burst frequency in DS excitatory neurons (Figure 4), establishing HCN1 as a tractable therapeutic target. Selective downregulation of HCN1, whether via small molecules or gene therapy, could offer a precision approach to reduce pathological bursting without broadly suppressing cortical activity. Moreover, the expression of HCN1 in the pacemaker cells of the heart ^70,71^ may also link this pathological overexpression to sudden unexpected death in epilepsy (SUDEP) ^72^ and other heart-related DS comorbidities ^73–75^. Indeed, HCN1 has been previously identified as a candidate gene for SUDEP ^76,77^ and overexpression of HCN channels in heart cells has been linked to ventricular arrhythmias in heart failure ^78–80^. Pharmacological intervention on HCN1 thus holds promise not only for seizure control but also for reducing systemic risk in DS patients.

In sum, our work highlights excitatory neuron dysfunction, via HCN1 overexpression, as a central, targetable feature of DS pathophysiology, and provides a strong rationale for the development of new therapies.

## Acknowledgments

We thank Jack Parent and Louis Dang for sharing the Dravet patient-derived hiPSC lines and Marcio Guiomar de Oliveira for deriving rat primary astrocytes. We also extend our thanks to the Advanced Cytometry Platform (Flow Core) at Guy’s Hospital and the Nikon Imaging Centre at King’s College London for their expertise and guidance in FACS and live calcium imaging experiments, respectively. Additionally, we thank Satinder Kaur Samra and the ARUK-UCL Drug Discovery Institute for helping and providing access to the MEA equipment and Fursham Hamid and Karla Lozano Gonzalez at the CDN-KCL Bioinformatics Core for performing the RNA-seq data analysis. We also thank Gabriele Lignani, for facilitating access to human postmortem tissue, Maria Thom, for locating and coordinating access to the tissue and the Epilepsy Society Brain and Tissue Bank, the Great Ormond Street Hospital Sample Bank and BRAIN UK at the University of Southampton for providing the necessary postmortem samples. HipSci Lines samples were collected from consented research volunteers recruited from the NIHR Cambridge BioResource through https://www.cambridgebioresource.group.cam.ac.uk/. The HipSci consortium obtained ethics approval for a revised consent (REC ref. 09/H0304/77, V3 15/03/2013), under which all data, except from the Y chromosome from males, can be made openly available (Y chromosome data can be used to de-identify men by surname matching). Finally, we thank Peter Harley and Mala Shah for comments on the manuscript. The authors acknowledge the Medical Research Council Center grant (MR/N026063/1). This research was funded, in part, by the Wellcome Trust. JB is supported by Wellcome Trust Investigator award (215508/Z/19/Z), UK Medical Research Council (MR/Z505353/1), the MND Association (2457-797) and funded through the generosity of a legacy donation from Margaret Potts. IL is supported by UK Medical Research Council (MR/Z505353/1), the MND Association (2457-797) and funded through the generosity of a legacy donation from Margaret Potts. JU is supported by UK Dementia Research Institute grant DRI-RE21605. FR is supported by Wellcome Trust “Cell Therapies & Regenerative Medicine” PhD studentship 108874/Z/15/Z.

---

**Figure S1.**
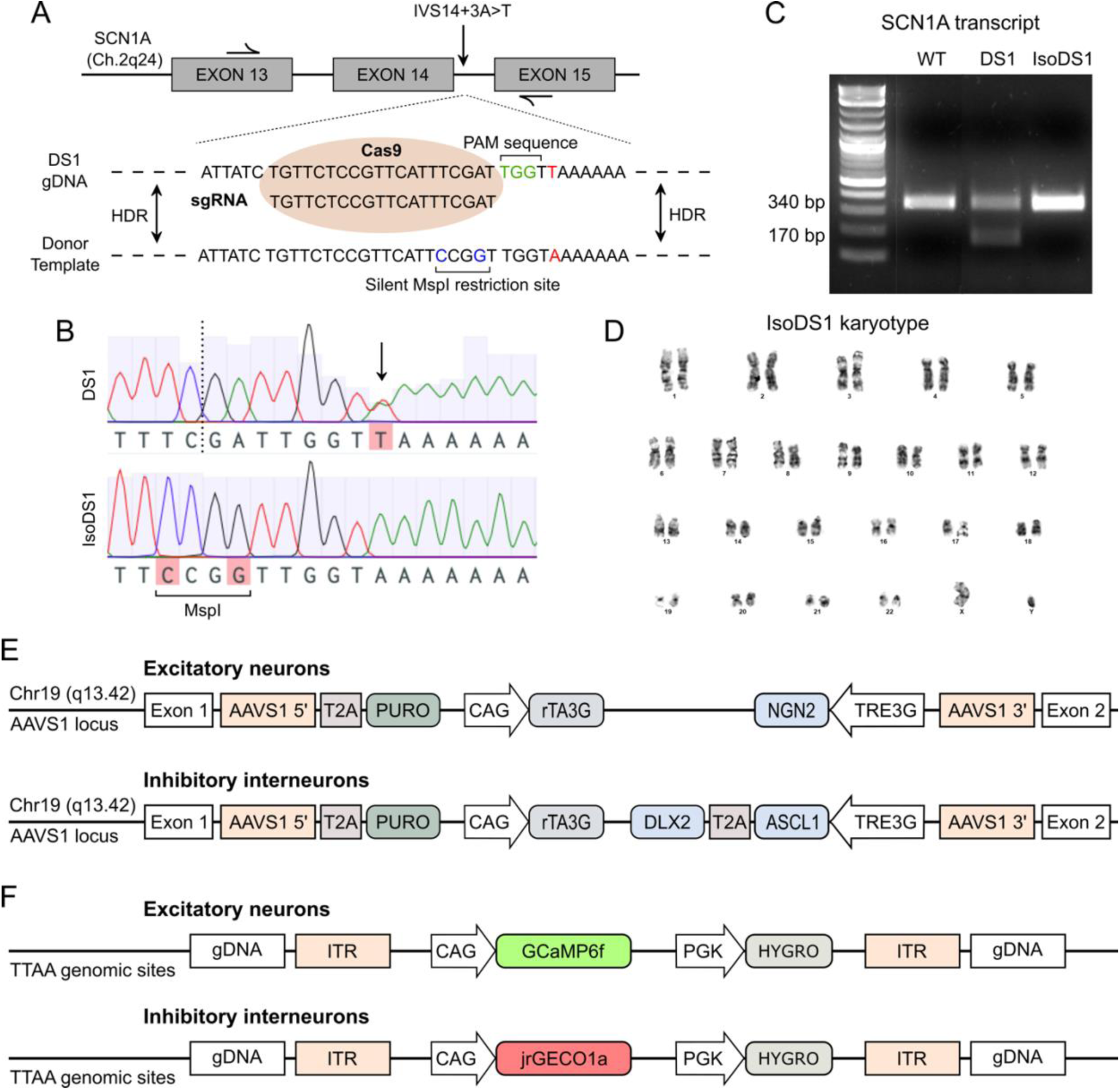
CRISPR/Cas9-mediated mutation correction and cell engineering. (**A**) Schematic showing the CRISPR/Cas9-mediated genome editing strategy to correct the endogenous *SCN1A*^IVS14+3A>T^ mutation in the DS1 line. Black arrows on exon 13 and 15 indicate the primers used for PCR amplification of the genomic region for sequencing purposes. Highlighted in red is the mutated nucleotide, in green the PAM sequence and in blue the silent mutations inserted in the donor template to introduce a MspI restriction site. (**B**) **–** Sanger sequencing of the genomic region of interest showing correction of the *SCN1A*^IVS14+3A>T^ mutation (black arrow) and homozygous integration of the donor template harbouring the silent MspI restriction site. The dotted line indicates the Cas9 cut site. (**C**) G-banding analysis showing normal karyotype for the DS1 corrected hiPSC line (IsoDS1). (**D**) Electrophoresis analysis of *SCN1A* transcript showing the full-size amplification of intron 13, exon 14 and intron 14 from WT cDNA (340 bp), the presence of 50% aberrant transcript missing exon 14 amplified from DS1 cDNA (170 bp) and the full-size amplification of the transcript fragment from corrected IsoDS1 cDNA (340 bp). (**E**) Schematic representation of the TALEN-mediated integration at the AAVS1 safe harbor locus of a vector containing, in between the 5’ and 3’ homology sequences, a puromycin resistance gene downstream of a T2A sequence, followed by a CAG promoter upstream of the rtTA3G. From the 3’ end, the TRE3G promoter regulates the doxycycline-inducible expression of the NGN2 transcription factor (excitatory neurons) or the ASCL1 and DLX2 transcription factors (inhibitory interneurons). (**F**) Schematic representation of the PiggyBac transposon-mediated integration, targeted at TTAA genomic sites, of a vector containing the calcium indicator downstream of a CAG promoter, followed by a hygromycin resistance cassette. The open reading frame is flanked by transposon-specific inverted terminal repeat sequences (ITRs).

**Figure S2.**
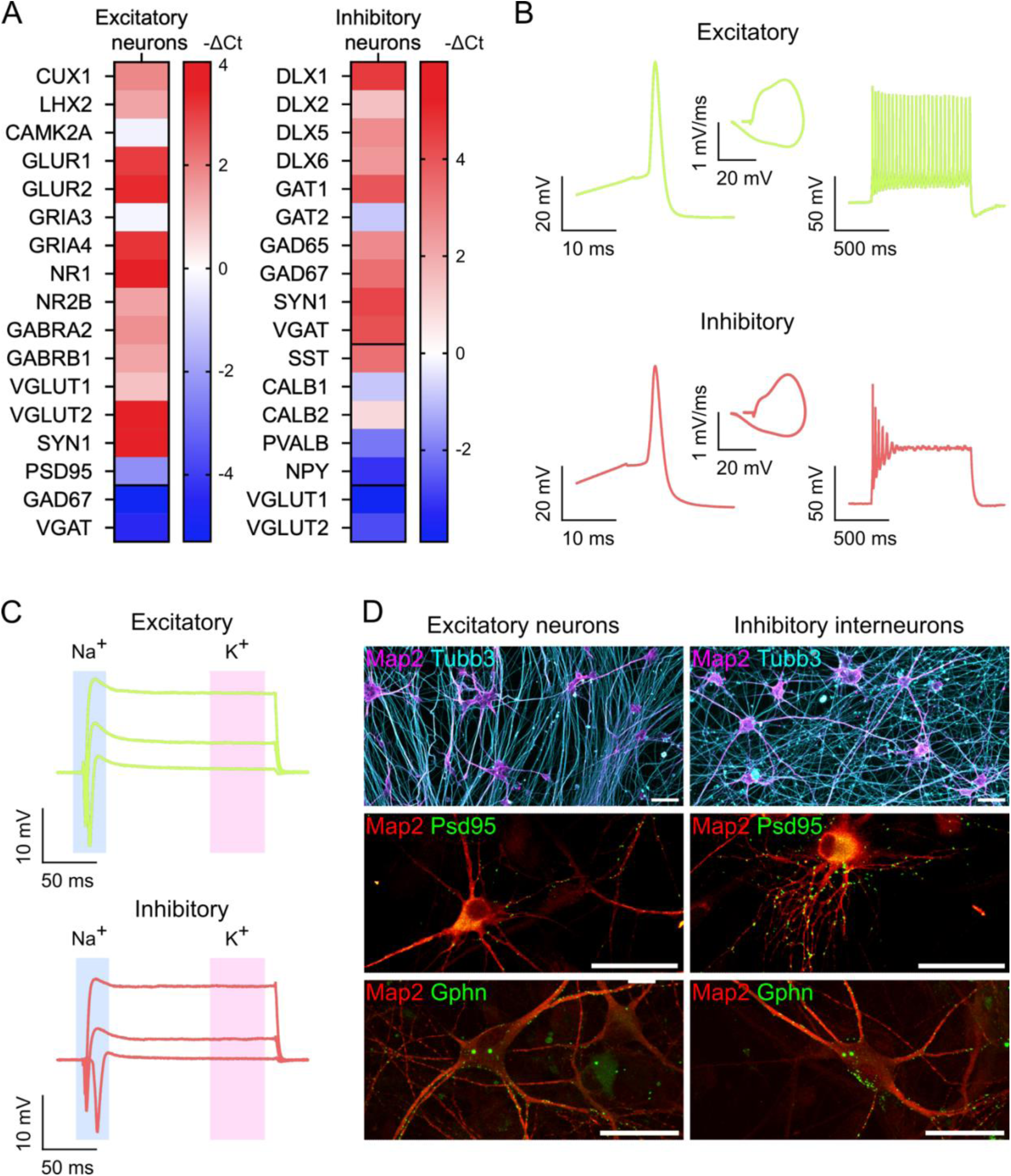
Characterisation of mature hiPSC-derived excitatory and inhibitory cortical neurons. (**A**) Heatmap representation of RT-qPCR analysis of wildtype excitatory and inhibitory neuron monocultures at D30. Gene expression levels are normalised to the housekeeping gene TBP and expressed as –ΔCt. n=3 independent experiments. (**B**) Representative single AP trace (left), phase plot (middle) and AP traces fired in response to 300pA of injected current (right) from wildtype excitatory and inhibitory neurons at 6 weeks. (**C**) Representative inward (Na^+^) and outward (K^+^) current traces recorded at –10, 30, and 70mV pulses from wildtype excitatory and inhibitory neurons at 6 weeks. (**D**) Top, immunofluorescence images of excitatory and inhibitory neurons stained for Map2 (magenta) and Tubb3 (cyan). Scale bar = 100μm. Bottom, immunofluorescence images of excitatory and inhibitory neurons stained for Map2 (red) and Psd95 or Gphn (green). Scale bar = 50μm.

**Figure S3.**
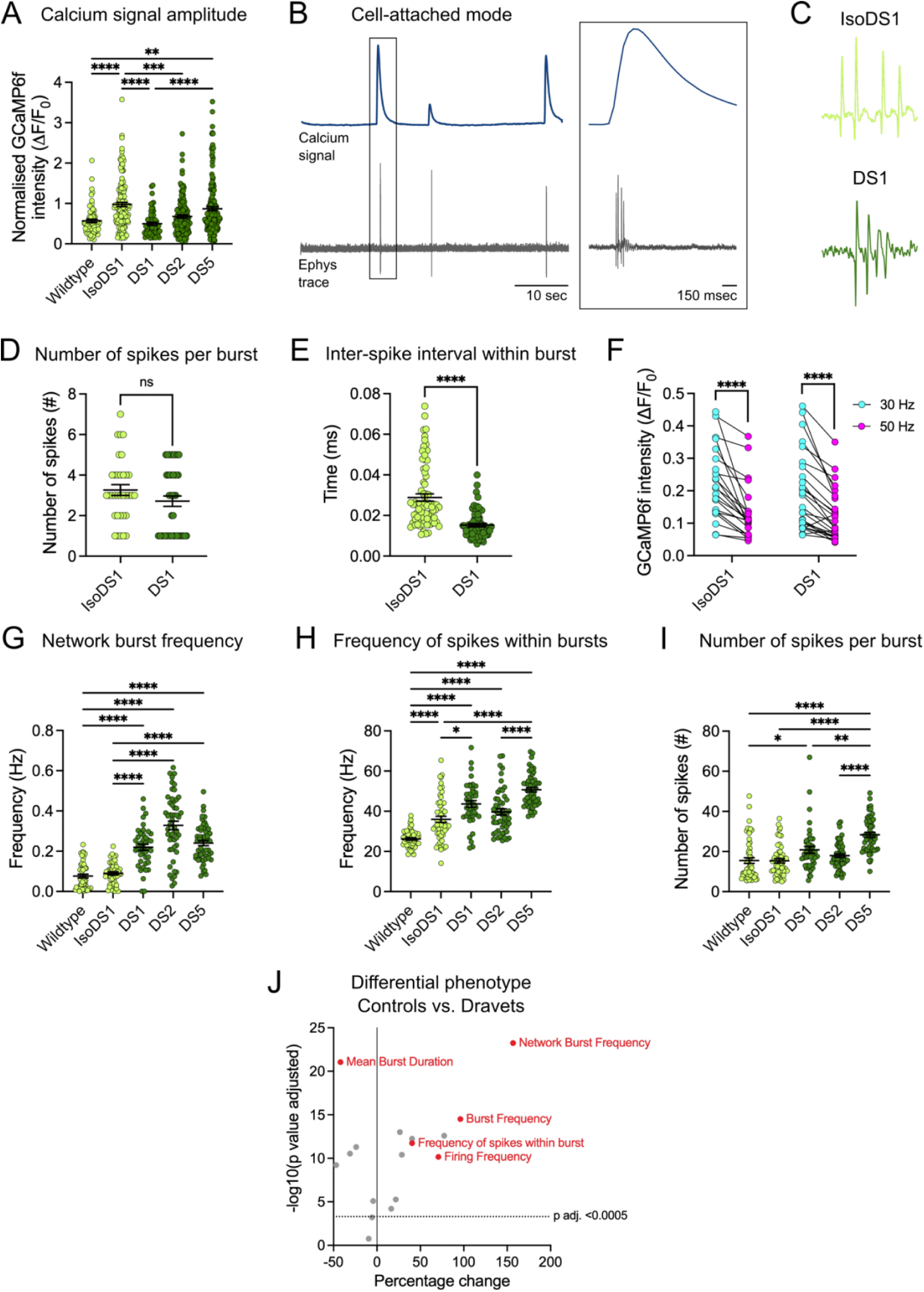
Increased neuronal activity of DS patient-derived excitatory and inhibitory co-culture networks. (**A**) Quantification of the single cell calcium signal amplitude (ΔF/F_0_) recorded for excitatory neurons in co-culture at 6 weeks (D50). Mean ± SEM, Kruskal-Wallis test with Dunn’s multiple comparison, ***p < 0.001, ****p < 0.0001, n=3/4 independent experiments (58-176 cells). (**B**) Example of a calcium signal trace (top, blue) and corresponding electrophysiological trace (bottom, grey) simultaneously recorded in cell-attached mode from a control excitatory neuron (IsoDS1). Box, detailed view of an individual calcium event and corresponding burst trace. (**C**) Representative burst traces recorded in cell-attached mode for IsoDS1 and DS1 excitatory neurons in co-culture. (**D-E**) Number of spikes per burst and inter-spike interval within burst quantified from cell-attached recordings in IsoDS1 and DS1 excitatory neurons at D50. Mean ± SEM, Mann-Whitney test, ns = not significant, ****p < 0.0001 (5-8 cells). (**F**) Quantification of the single cell calcium signal amplitude recorded for IsoDS1 and DS1 excitatory neurons in response to 30 and 50 Hz field stimulation. Wilcoxon matched-pairs signed rank test, ****p < 0.0001 (23-26 cells). (**G-I**) Quantification of network burst frequency, frequency of spikes within burst and number of spikes per burst recorded using MEAs during week 6 of co-culture. Mean ± SEM, Kruskal-Wallis test with Dunn’s multiple comparison, *p < 0.05, **p < 0.01, ****p < 0.0001, n=3 independent experiments (each with 6 wells per genotype, 3 recordings – D42, D46, D50). (**J**) Volcano plot showing differential phenotypical characteristics in Dravet (DS1, DS2 and DS5) vs. control (wildtype and IsoDS1) co-cultures based on electrophysiological parameters (listed in the method section) extrapolated from field potential recordings (MEAs) at week 6 (D42, D46, D50). Highlighted in red are some of the most significant phenotypic changes.

**Figure S4.**
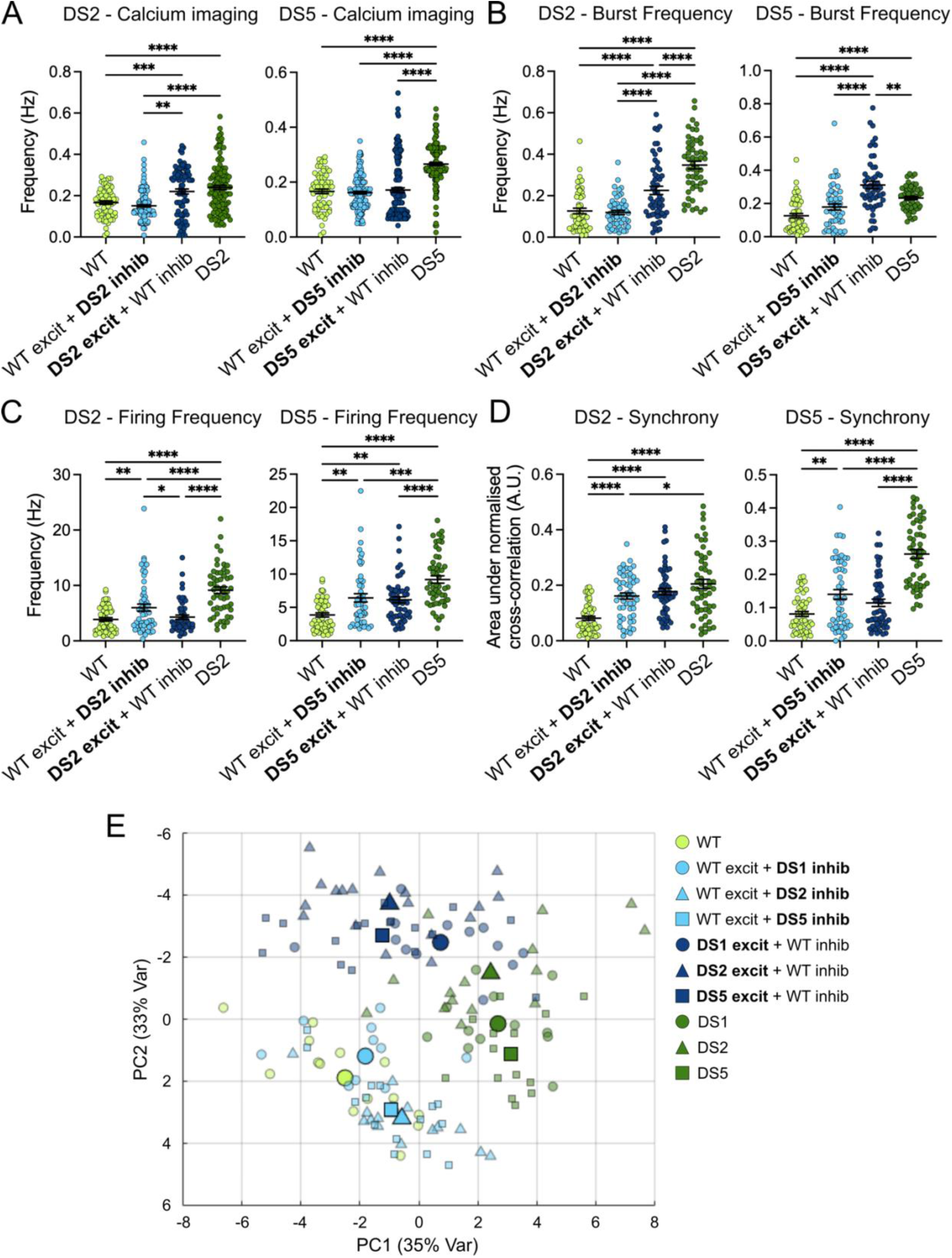
Changes in neuronal activity in mixed-genotype excitatory/inhibitory co-cultures. (**A**) Quantification of calcium signal frequency recorded for excitatory neurons (GCaMP6f) in DS2 and DS5 mixed– and same-genotype co-cultures at 6 weeks (D50). Mean ± SEM, Two-way ANOVA with Tukey’s multiple comparison, *p < 0.05, ****p < 0.0001, n=3 independent experiments (43-176 cells). (**B-D**) Quantification of the burst frequency, firing frequency and synchrony recorded in the different mixed– and same-genotype co-culture conditions during week 6. Mean ± SEM, Two-way ANOVA with Tukey’s multiple comparison, *p < 0.05, **p < 0.01, ***p < 0.001, ****p < 0.0001, n=3 independent experiments (each with 6 wells per genotype, 3 recordings – D42, D46, D50). (**E**) PCA of electrophysiological parameters, listed in the methods section, extrapolated from field potential recordings (MEAs) of mixed– and same-genotype co-cultures at week 6 (D42, D46, D50). Smaller symbols represent individual wells, bigger symbols represent the average per condition.

**Figure S5.**
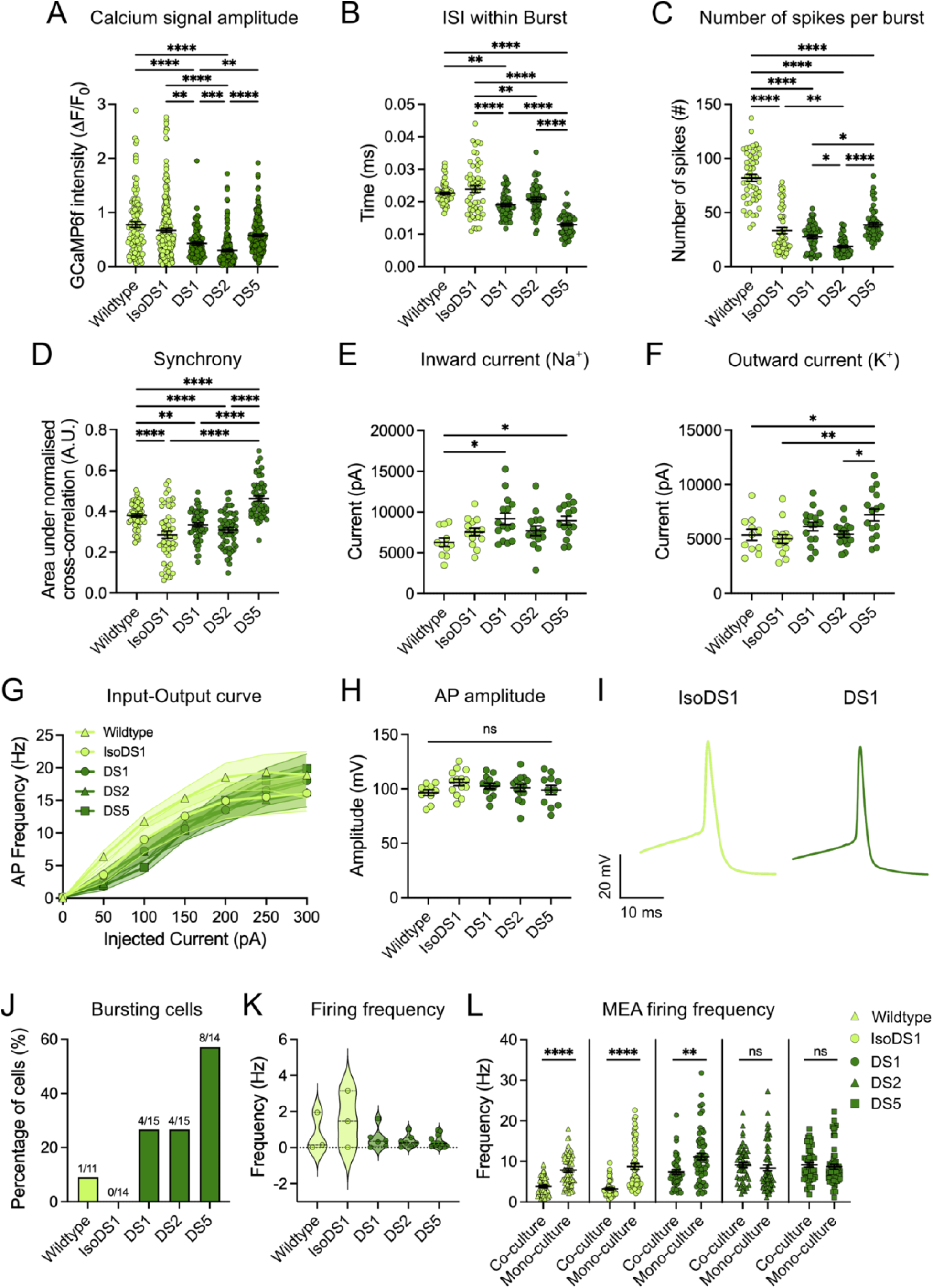
Characterisation of neuronal activity of DS patient-derived excitatory neurons in monoculture. (**A**) Quantification of the single cell calcium signal amplitude (ΔF/F_0_) recorded for excitatory neurons in monoculture at 6 weeks (D50). Mean ± SEM, Kruskal-Wallis test with Dunn’s multiple comparison, ***p < 0.001, ****p < 0.0001, n=3/4 independent experiments (60-135 cells). (**B-D**) Inter-spike interval within burst, number of spikes per burst and synchrony quantified from field potential recordings of excitatory neurons in monoculture at week 6. Mean ± SEM, Kruskal-Wallis test with Dunn’s multiple comparison, *p < 0.05, **p < 0.01, ****p < 0.0001, n=3 independent experiments (each with 6 wells per genotype, 3 recordings – D42, D46, D50). (**E-F**) Quantification of inward (Na^+^) and outward (K^+^) currents recorded in excitatory neurons at 6 weeks. Mean ± SEM, One-way ANOVA with Tukey’s multiple comparison, *p < 0.05, **p < 0.01, n=3 independent experiments (11-15 cells). (**G**) AP frequency in relation to injected current recorded in excitatory neuron monocultures at 6 weeks. Mean ± SEM. n=3 independent experiments (11-15 cells). (**H**) Quantification of AP amplitude recorded in excitatory neurons at 6 weeks. Mean ± SEM, One-way ANOVA with Tukey’s multiple comparison, ns = not significant, n=3 independent experiments (11-15 cells). (**I**) Representative single AP traces from IsoDS1 and DS1 excitatory neurons at 6 weeks. (**J**) Quantification of the percentage of bursting excitatory neurons per condition. n=3 independent experiments (11-15 cells). (**K**) Firing frequency from spontaneous current-clamp recordings, quantified only for active cells. n=3 independent experiments (3-12 cells). (**L**) Quantification of firing frequency from field potential recordings (MEAs) in co-culture compared to monoculture conditions at 6 weeks. Mean ± SEM, Mann-Whitney test, ns = not significant, **p < 0.01, ****p < 0.0001, n=3 independent experiments (each with 6 wells per genotype, 3 recordings – D42, D46, D50).

**Figure S6.**
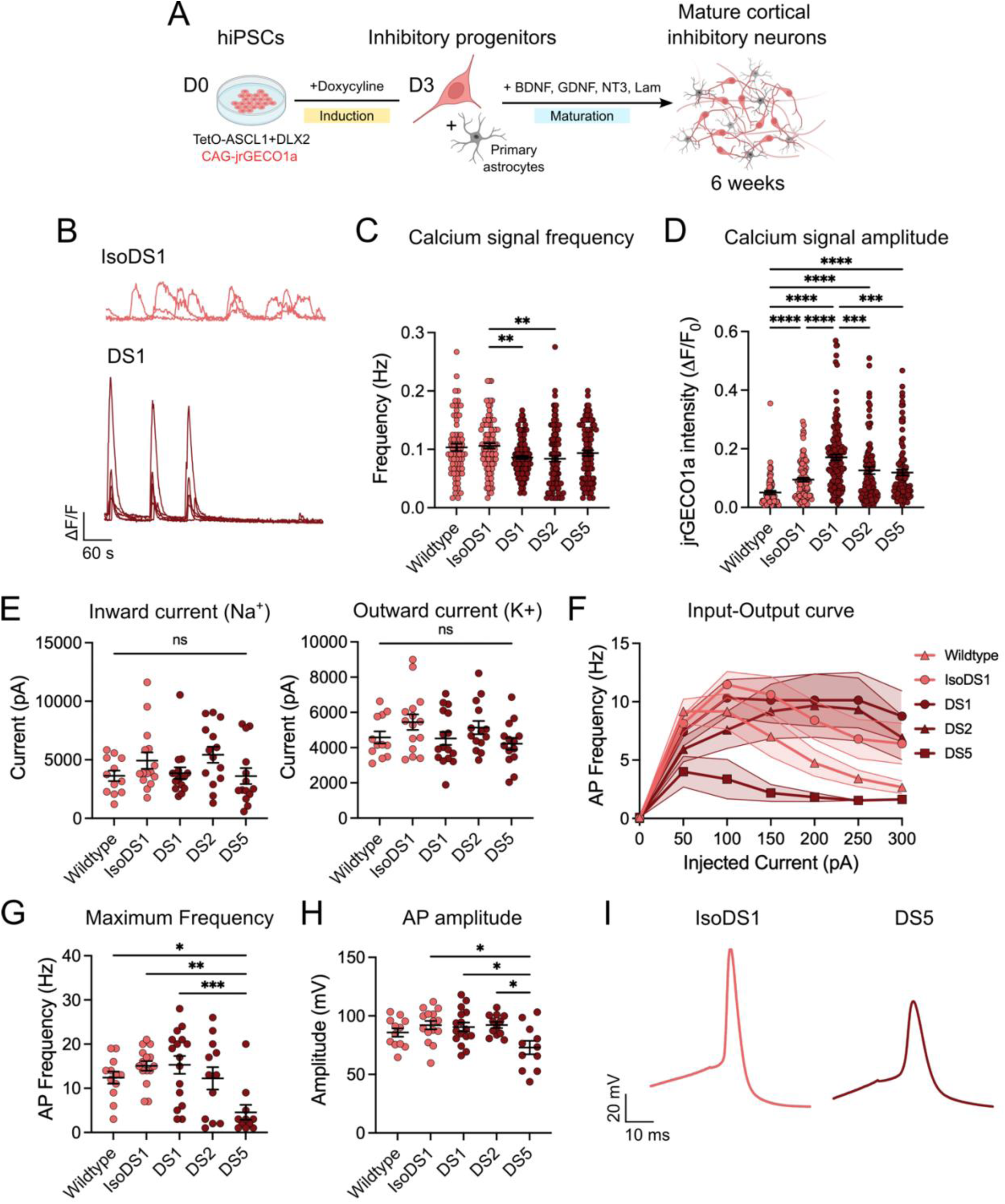
Characterisation of neuronal activity of DS patient-derived inhibitory interneurons in monoculture. (**A**) A schematic of the forward programming differentiation into mature cortical inhibitory interneurons. (**B**) Representative calcium traces from IsoDS1 and DS1 interneuron monocultures at 6 weeks (D50), plotted as the ΔF/F_0_ over time (60 sec). (**C-D**) Quantification of calcium signal frequency and amplitude recorded from interneuron monocultures at 6 weeks (D50). Mean ± SEM, Kruskal-Wallis test with Dunn’s multiple comparison, **p < 0.01, ***p < 0.001, ****p < 0.0001, n=3 independent experiments (70-132 cells). (**E**) Quantification of inward (Na^+^) and outward (K^+^) currents recorded in inhibitory interneurons at 6 weeks. Mean ± SEM, Kruskal-Wallis test with Dunn’s multiple comparison and One-way ANOVA with Tukey’s multiple comparison, ns = not significant, n=3 independent experiments (12-16 cells). (**F**) AP frequency in relation to injected current recorded in interneuron monocultures at 6 weeks. Mean ± SEM. n=3 independent experiments (12-16 cells). (**G-H**) Quantification of the maximum evoked AP frequency and AP amplitude recorded in interneuron monocultures at 6 weeks. Mean ± SEM, One-way ANOVA with Tukey’s multiple comparison, *p < 0.05, **p < 0.01, ***p < 0.001, n=3 independent experiments (12-16 cells). (**I**) Representative single AP traces from IsoDS1 and DS5 inhibitory interneurons at 6 weeks.

**Figure S7.**
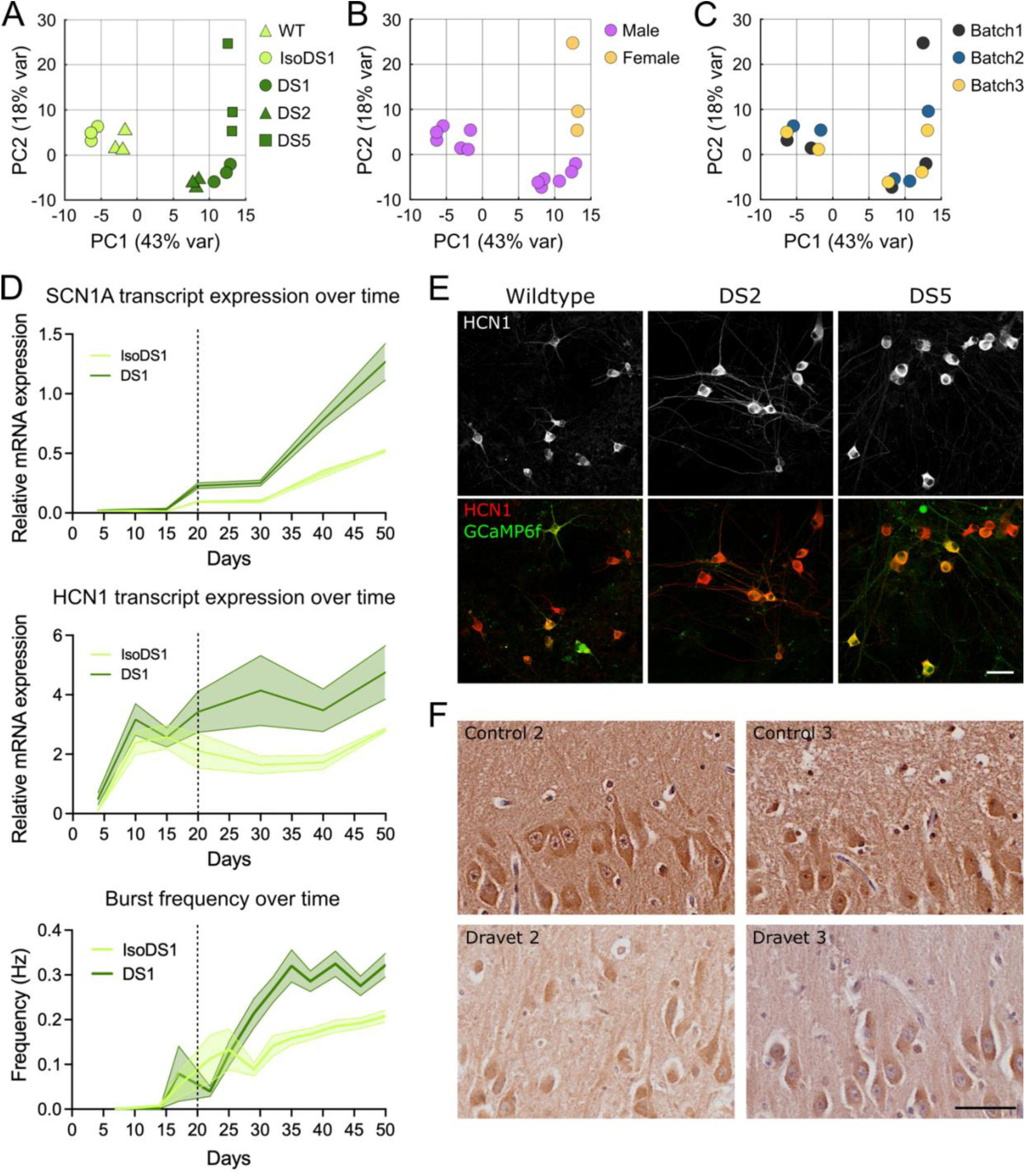
HCN1 expression in DS patient-derived excitatory neurons. (**A**-**C**) PCA of bulk RNA-seq data from excitatory neurons at 6 weeks (D50) showing data clustering by genotype, sex and by sample processing batch. (**D**) Quantification of SCN1A (top) and HCN1 (middle) transcript expression over time in IsoDS1 and DS1 excitatory neurons in relation to the appearance of the burst frequency phenotype (bottom). The dotted line corresponds to the time-point when SCN1A is first detected. Mean ± SEM, n=3 independent experiments. (**E**) Fluorescence images of Wildtype, DS2 and DS5 excitatory neurons at 6 weeks (D50) stained for HCN1 (red), GCaMP6f (green) and TUBB3. Scale bar 50 µm. (**F**) Representative images of human hippocampal slices from control and Dravet syndrome cases stained for HCN1. Scale bar 50 µm.

**Figure S8.**
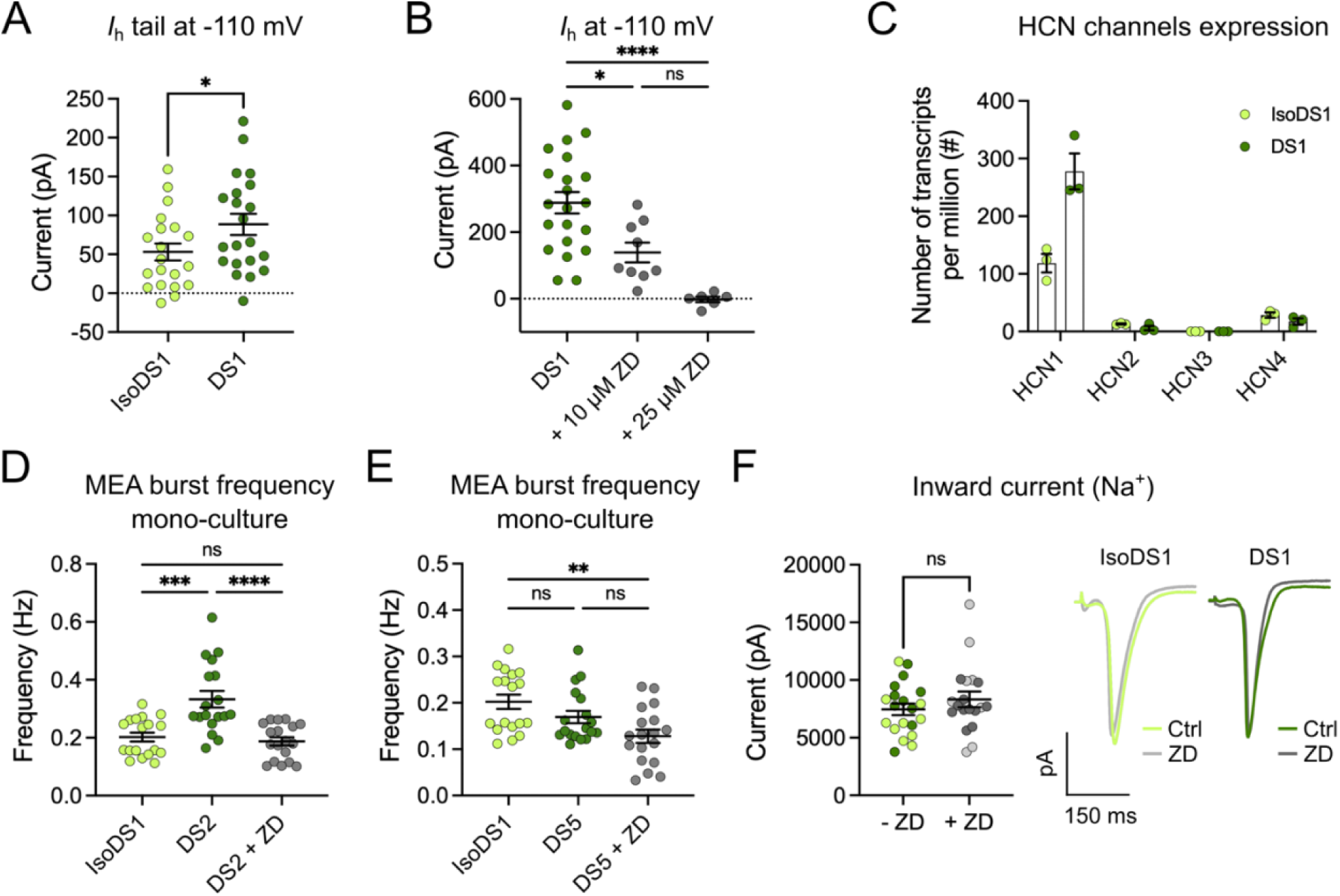
H current in DS patient-derived excitatory neurons. (**A**) Quantification of *I*_h_ tail current amplitude at the last voltage step (−110mV) in IsoDS1 and DS1 excitatory neurons at 6 weeks (D50). Mean ± SEM, Welch’s t-test, *p < 0.05 (19-21 cells). (**B**) Quantification of *I*_h_ amplitude at the last voltage step (−110mV) in DS1 excitatory neurons treated with either 10µM or 25 µM ZD 7288 at 6 weeks (D50). Mean ± SEM, One-way ANOVA with Tukey’s multiple comparison, ns = not significant, *p < 0.05, ****p < 0.001 (6-21 cells). (**C**) HCN1, HCN2, HCN3 and HCN4 transcript count from bulk RNA-seq in IsoDS1 and DS1 excitatory neurons at 6 weeks (D50). Mean ± SEM, n=3 independent experiments. (**D**-**E**) Quantification of the burst frequency recorded for IsoDS1 and DS2 or DS5 excitatory neurons as well as DS2 or DS5 excitatory neurons treated with 25 µM ZD 7288 at D50. Mean ± SEM, Kruskal-Wallis test with Dunn’s multiple comparison, ns = not significant (p>0.05), **p < 0.01, ***p < 0.001, ****p < 0.0001, n=3 independent experiments (each with 6 wells per genotype). (**F**) Left, quantification of Na^+^ currents recorded at 6 weeks in DS1 (dark green/dark grey) and IsoDS1 (light green/light grey) excitatory neurons either untreated or treated with 25 µM ZD 7288. Mean ± SEM, Mann-Whitney test, ns = not significant (19-20 cells). Right, representative inward Na^+^ current traces recorded at 10mV from IsoDS1 and DS1 excitatory neurons untreated or treated with 25 µM ZD 7288.

## MATERIALS AND METHODS

### Molecular cloning and plasmid generation

Plasmid pSpCas9(BB)-2A-Puro (PX459) V2.0 (Addgene #62988) was digested with BbsI enzyme and ligated with annealed primers designed to contain the entire sgRNA sequence necessary for CRISPR-correction of the DS1 line. Plasmid pUCM-AAVS1-TO-hNGN2 (Addgene #105840) was digested with PmeI and KpnI to remove the fragment ‘*EF1a-mCherry’* and maintain the *TRE3G-hNGN2_CAG-rTA3G* cassette. The resulting plasmid was digested with PacI and NotI to replace hNGN2 with a PCR-derived fragment containing hASCLI and hDLX2 inserted either side of a T2A self-cleaving peptide to generate the cassette *TRE3G-hASCL1-T2A-hDLX2_CAG-rTA3G*. hASCL1 and hDLX2 were subcloned from plasmid phASCL1-N106 (Addgene #31781) and phDLX2-N174 (Addgene #60860), respectively. GCaMP6f and jRGECO1a were PCR-amplified from plasmids pGP-CMV-GCaMP6f (Addgene #40755) and pGP-CMV-NES-jRGECO1a (Addgene #61563), respectively, and inserted downstream of a CAG promoter into a PiggyBAC backbone ^81^.

Cloning was performed by a combination of enzymatic digestion (NEB) and PCR amplification using a Q5 polymerase system (NEB) and Gibson Assembly (NEB). All primers used are listed in Table S1.

### CRISPR/Cas9-mediated correction of the DS1 cell line

Targeted gene editing of the DS1 line to correct the endogenous *SCN1A*^IVS^ ^14+3A>T^ mutation (genome coordinates NC_000002.11[166,039,420]) was achieved using CRISPR/Cas9-mediated HDR. The sgRNA (CRISPR ID: 941482857, Table S2) was designed using the Wellcome Sanger Institute Genome Editing (WGE) online tool, chosen based on its proximity to the mutation and its off-target score and cloned into Addgene plasmid #62988 as described above. The double-strand oligonucleotide donor template (Sigma-Aldrich, Table S2) was designed according to the wildtype sequence and it was modified to introduce a silent MspI restriction site in the seed region of the sgRNA binding site of the CRISPR edited cells. The plasmid expressing the sgRNA, spCas9 and a puromycin resistance gene, was delivered by electroporation together with the donor template and 24h later cells were treated with puromycin (0.5 μg/mL) for 24h. Single-cell clones were derived and screened for the presence of the MspI restriction site by PCR and enzymatic digestion using Taq 2X Master Mix (NEB), primers SCN1A-14/13_2F and SCN1A-14/13_3R (Table S2) and MspI restriction enzyme (NEB). Positive clones were Sanger sequenced (SourceBioscience) to confirm successful correction as well as the integrity of the genomic sequence around the editing site (∼1000 bp upstream and downstream of the mutation). The top 5 off-target sites were also sequenced to confirm the absence of off-target editing (Table S2). Finally, g-banding was performed to confirm a normal karyotype (CellGuidanceSystems).

### Human induced Pluripotent Stem Cell (hiPSC) engineering

Patient-derived hiPSC lines harboring the pathogenic *SCN1A*^IVS14+3A>T^ (DS1), *SCN1A*^Y325X^ (DS2) ^12^ and *SCN1A*^R222X^ (DS5) ^17^ mutations were provided by Dr Louis Dang and Professor Jack Parent (University of Michigan). The IsoDS1 line (isogenic control for the DS1 line) was generated in house as described above. The wildtype PAMV_1 line was obtained from the human induced pluripotent stem cell initiative (HiPSCi).

hiPSCs intended for forward programming differentiation into excitatory or inhibitory neurons were engineered to express the *TRE3G-hNGN2_CAG-rTA3G* or *TRE3G-hASCL1-T2A-hDLX2_CAG-rTA3G* cassette, respectively, using TALEN-mediated integration into the AAVS1 safe-harbour locus (fig. S1E). Donor plasmid and TALENs (Addgene #52341 and #52342) were delivered by electroporation, positively transfected cells were enriched by puromycin selection (0.5 μg/mL) and homozygous clones were identified by PCR screening as described by Fernandopulle et al. ^18^. Engineered hiPSC clones were further modified to express either the genetically encoded calcium indicator *CAG-GCaMP6f* or *CAG-jrGECO1a* using PiggyBAC-mediated transposon integration (fig. S1F). Donor plasmid and transposase were delivered by electroporation and positive cells were isolated using fluorescence-activated cell sorting (FACS).

### hiPSC culture and differentiation

hiPSCs were maintained on 3.5 μg/ml LN521-coated plates (BioLamina) in StemMACS iPS-Brew XF (Miltenyi Biotec). Cells were passaged at 70% confluency using TrypLE Express (Gibco) and replated in medium supplemented with 10 μM Y-27632 (Tocris) for 24h.

Excitatory and inhibitory neuron differentiation was based on previously published protocols ^18,19,82^ with minor modifications. For **excitatory neuron differentiation**, on day 0 (D0), 70% confluent hiPSCs were dissociated with TrypLE Express, resuspended in Induction Medium (Table S3) supplemented with 10 µM ROCK inhibitor Y-27632 (Tocris) and plated at 100.000 cells/cm^2^ into dishes coated with 200 μg/ml GFR Matrigel (Corning). After 24h, Y-27632 was removed and Induction Medium was replaced daily until Day 3 (D3), when the progenitors were dissociated with Accumax (Millipore) and either frozen or plated for maturation. For **inhibitory interneuron differentiation**, 70% confluent hiPSCs were dissociated with TrypLE Express, resuspended in StemMACS iPS-Brew XF supplemented with 10 µM Y-27632 and plated at 40.000 cells/cm^2^ onto GFR Matrigel-coated dishes. After 24h, Y-27632 was removed. After a further 24h, on Day 0 (D0), StemMACS iPS-Brew XF medium was replaced to Induction Medium, which was replaced daily until Day 4 (D4), when the progenitors were dissociated with Accumax (Millipore) and either frozen or plated for maturation.

After pre-differentiation, excitatory and inhibitory progenitors were resuspended in Cortical Maturation Medium (Table S3) supplemented with 10µM Y-27632 and plated onto a monolayer of rat primary astrocytes for maturation. Seeding densities are specified below for each technique. 24h post-plating, half the culture medium was replaced by an equal volume of fresh Cortical Maturation Medium supplemented with 10µM Ara-C (final concentration 5µM) (Sigma). From Day 7 onwards, half medium changes were performed once a week.

### Immunocytochemistry and image analysis

For imaging experiments, neuronal progenitors were plated at the density of 50.000 cells/cm^2^ on a monolayer of primary astrocytes previously plated on GFR-Matrigel-coated imaging plates. For co-cultures, seeding densities comprised 70% excitatory neurons and 30% inhibitory interneurons. At the desired time point, samples were fixed with 4% PFA (Thermo Fisher Scientific) for 10 min, washed 3x with DPBS, and permeabilised and blocked for 30 min with 3% BSA (Sigma-Aldrich), 0.1% Triton X-100 (Thermo Fisher Scientific) in DPBS. Primary antibodies (Table S4), appropriately diluted in block buffer, were applied overnight at 4°C. Samples were washed 3x with DPBS before secondary antibodies (Table S4) were applied for 1h at RT. Following the final three DPBS washes, samples were imaged using the confocal laser scanning microscope Leica TCS SP8 (Leica Microsystems).

Image processing was performed using FiJi (NIH) and a custom-made macro deposited at https://github.com/jburrone. For excitatory/inhibitory cell type ratio analysis, for each field of view GFP^+^ and RFP^+^ neurons were manually counted and ratios were calculated in Microsoft Excel. For HCN1 fluorescence intensity quantification, each field of view was analysed as follow: the TUBB3 signal (647 channel) was used to generate a mask of **total neuronal area** by applying Gaussian Blur, Auto Threshold and Remove Outliers. The “total neuronal area mask” was then overlayed onto the HCN1 signal (568 channel) and the Mean Gray Value within the mask was measured to obtain HCN1 fluorescence intensity in the neurons; the GCaMP6f signal (488 channel) was used to generate a mask of **somatic area** by applying Gaussian Blur, Auto Threshold and Analyse Particles. The “soma mask” was then overlayed onto the HCN1 signal (568 channel) and the Mean Gray Value within the mask was measured to obtain HCN1 fluorescence intensity in the somas; finally, the “neuronal bodies mask” was subtracted from the “total neuronal area mask” to obtain a mask of **neuronal processes area**. The “neuronal processes mask” was then overlayed onto the HCN1 signal (568 channel) and the Mean Gray Value within the mask was measured to obtain HCN1 fluorescence intensity in the dendrites. Finally, the HCN1 dendrite/soma ratio were manually calculated in Microsoft Excel.

### Immunohistochemistry and image analysis of human tissue

Formalin-fixed paraffin-embedded 6 µm sections from 3 control and 3 Dravet patients hippocampus were received from the Great Ormond Street Hospital Sample Bank, Brain UK at the University of Southampton and the Epilepsy Society Brain and Tissue Bank. After the sections were baked overnight at 60°C, they were taken through xylene for de-waxing and underwent rehydration through a series of ethanol solutions (100%, 90% and 70%). Endogenous peroxidase activity was blocked with incubation in methanol and 3% hydrogen peroxide solution for 10 minutes. Heat-mediated epitope retrieval was performed with Tris-EDTA pH 9 in a pressure cooker for 10 minutes. Sections were blocked in 10% milk solution for 30 minutes before being incubated with primary antibody (HCN1 – Proteintech, 55222-1-AP, 1:100) overnight at 4°C. Sections were washed three times in TBS-Tween before secondary antibody (Goat anti-rabbit Biotinylated – Vector BA-1000, 1:200) was applied for 30 minutes. Sections were washed three times in TBS-Tween and then incubated in Avitin-Biotin Complex (Vectastain ABC Kit Peroxidase, PK-4000) for 30 minutes, washed again and then incubated for 4 minutes in DAB substrate activated with hydrogen peroxide. Nuclei were counterstained in Mayer’s haematoxylin and then dehydrated via a series of increasing ethanol solutions. Sections were incubated in xylene and mounted using DPX non-aqueous mounting medium. Finally, slides were then scanned at 20x magnification using Olympus VS120 digital slide scanner.

One equal size field of view was extracted from each scanned image using the QuPath software. Further image processing was performed using FiJi (NIH) and a custom-made macro deposited at https://github.com/jburrone. For the quantification of DAB^+^ dendritic area, each field of view was analysed as follow: the DAB channel was used to generate a dendritic mask by applying MorphoLibJ Directional Filtering, Ridge Detection, MorphoLibJ Analyse Regions and filtering based on Oriented Box Orientation between 45 and 135 degrees. The obtained mask was then dilated and used to measure the area occupied by DAB^+^ dendrites.

### RT-PCR and RT-qPCR

For RT-q/PCR, neuronal progenitors were plated at the density of 170.000 cells/cm^2^ on a monolayer of primary astrocytes previously plated on GFR-Matrigel-coated plates. At the desired time point, total RNA was isolated using RNeasy Mini Kit (Qiagen) and first-strand cDNA was obtained from retro transcription of RNA using GoScript^TM^ Reverse Transcription System (Promega).

For *SCN1A* transcript analysis (fig. S1C), cDNA from 3-week-old wildtype, DS1 and IsoDS1 excitatory neurons was used as a template for PCR amplification using Q5 High-Fidelity 2X Master Mix (NEB) and SCN1A_13/15 forward and reverse primers (Sigma-Aldrich – Table S5) to amplify the portion of the transcript comprised between exon 13 and 15. The resulting PCR products were run on a 1.5% agarose gel and the gel was imaged using a Bio-Rad Gel Dock XR+ Molecular Imager (Bio-Rad).

For neuronal gene expression characterisation, quantitative RT-PCR was carried out in 10μl reaction mixtures containing 10-20 ng cDNA derived from 4-week-old neuronal cultures, oligonucleotide primers (Sigma-Aldrich – Table S5) and Fast SYBR Green PCR Master Mix (Thermo Fischer Scientific). RT-qPCR was performed using a CFX384 Touch Real-Time PCR Detection System (Bio-Rad). For gene expression analysis, Ct values of target genes were normalised to the Ct value of the housekeeping gene TBP. Normalised data was displayed as a – ΔCt heatmap. For *SCN1A* and *HCN1* expression time-course, data was analysed by Comparative CT Method ^83^.

### Whole-cell patch clamp electrophysiology

For patch clamp recordings, neuronal progenitors were plated at the density of 40.000 cells/cm^2^ on a monolayer of primary astrocytes previously plated on GFR matrigel-coated 18mm glass coverslips. For co-cultures, seeding densities comprised 70% excitatory neurons and 30% inhibitory interneurons. Excitatory and inhibitory neurons in co-culture were distinguished by the presence of either GCaMP6f (detected using a 488 nm filter) or jrGECO1a (detected using a 568 nm filter), respectively.

After 6 weeks of maturation, coverslips were transferred to an open bath chamber (Warner Instruments) containing extracellular solution (Table S6) at pH 7.3 and osmolarity at ∼305 mOsm. For recordings in the presence of the H current blocker ZD 7288, either 10 or 25 µM ZD 7288 (Tocris) were added to the extracellular solution. The chamber was mounted on an inverted epifluorescent microscope (Olympus IX71) fitted with a 60x oil objective. Glass pipettes were pulled from borosilicate glass (O.D. 1.5 mm, I.D. 0.86 mm, Sutter Instruments) to have a resistance between 3 and 5 MΩ and were filled with 10 µl of intracellular solution (Table S6) at pH 7.4 and osmolarity 290 mOsm. Whole-cell patch clamp was performed at the soma of neurons using a Multiclamp 700B amplifier (Molecular Devices) and recorded using a Digidata 1440A digitizer (Molecular Devices). All recordings were carried out at room temperature and the data was acquired with Clampex software (Molecular Devices) and Axon Multiclamp Commander Software (Molecular Devices). Current-clamp data was sampled at a rate of 50 kHz and filtered at 10 kHz while voltage-clamp data was sampled at 20 kHz and filtered at 10 kHz. Whole-cell currents used to estimate the conductance of Na^+^ and K^+^ were recorded in voltage-clamp mode using 50 msec voltage steps from –80 mV to +50 mV. Values were corrected for baseline current offset before stimulation. H currents were also recorded in voltage-clamp mode, using 1 sec voltage steps from –60 mV to –110 mV (10 pA steps). Resting membrane potential and spontaneous action potential (AP) spiking were recorded, within the first 2 minutes after whole-cell patch was achieved, for 1 minute in current-clamp mode in the absence of current injection. Intrinsic excitability measurements and AP properties were recorded in current-clamp mode, using a steady current injection to maintain the membrane potential at –60 mV. Input-Output measurements were performed using 500 msec current injections from –50 pA to 300 pA (50 pA steps) while AP properties were assessed using 10 msec current injections from –20pA to 170pA (10 pA steps).

Electrophysiological measurements were analysed using custom MATLAB scripts. Inward Na^+^ current amplitudes were measured as the minimum value of a current trace, while steady state outward K^+^ current amplitudes were measured by averaging values over a 15 msec window acquired 25 msec after the voltage step. H current amplitudes were measured as the delta between averaged values the end of the voltage step (200 msec window) and averaged values at the beginning of the voltage step (20 msec window).

AP properties were measured using sequential injection of 10 msec current steps of increasing amplitude (10 pA increments). Only the measurements of the first AP at the current threshold (first step to elicit an AP) were considered. AP waveforms were extracted using the MATLAB’s findpeaks function with minimum peak Amplitude 0 mV. For each cell we extracted the amplitude (max amplitude – average Vm at the end of stimulus 50 msec window excluding APs) and voltage threshold (voltage at the time the speed of Vm rise is above 0.15 mV/ms). Input-Output curves were obtained using sequential injection of 500 msec current steps of increasing amplitude (50 pA increments). The location of the AP was extracted using MATLAB’s findpeaks function with minimum peak Amplitude set at 0 mV. Cells with access resistance greater than 30 MΩ or a holding current lower than –100 pA were excluded from the analysis.

### Calcium imaging

For calcium imaging experiments, neuronal progenitors were plated at the density of 50.000 cells/cm^2^ on a monolayer of primary astrocytes previously plated on GFR-Matrigel-coated imaging plates. For co-cultures, seeding densities comprised 70% excitatory neurons and 30% inhibitory interneurons.

At the time of imaging, neurons were gently washed in warm HBS (Table S7) before warm Ca^2+^ Imaging Buffer (Table S7) was added. After 1-2 min equilibration, Ca^2+^ imaging was performed at room temperature using a Nikon Eclipse Ti-2 inverted microscope equipped with 2 x Photometrics Prime 95B sCMOS cameras. GCaMP6f fluorescence was recorder using a 488 nm filter while jrGECO1a fluorescence was recorder using a 568 nm filter. Time-lapse recordings were acquired using a 20x objective, at a 1x digital zoom, for 2 minutes at a frame rate of 10-20 frames/sec. For 2-colour imaging, the dual-camera mode was used and time-lapse recordings were acquired using a 20x objective – 1x digital zoom, with 488-568 sequential/alternate frame acquisition at a frame rate of 9 frames/sec. An average of 2-3 fields of view per coverslip were imaged and a minimum of 3 coverslip per biological sample were included in the analysis.

Time-lapses were analysed using a series of published MATLAB script ^84^. The script “mmc3_REVISED” was used to detect ROIs and calculate the raw fluorescent intensity over time for each ROI. The script “mmc6” was used to detect calcium spikes and calculate the spike rate and average spike amplitude for each ROI, as well as to calculate the normalised fluorescence intensity over time (ΔF/F_0_) for each ROI.

### Multi Electrode Array (MEA) recording

For MEA recordings, CytoView MEA 96 well plates (Axion Biosystems) were first coated in a drop-wise manner (8 µl) with PLO and mouse Laminin (ThermoFisher). An 8 µl mix of 50.000 neuronal progenitors and 12.500 primary astrocytes was then plated in the centre of each well and the cells were left to adhere in the incubator for 1-2 h before the total volume of media was added.

For co-cultures, neuronal densities comprised 70% excitatory neurons and 30% inhibitory interneurons. Starting from Day 7, neuronal activity was recorded for 15 minutes every 3-4 days until Day 50 using a Maestro Pro microelectrode array (Neural Real-Time – Spontaneous; Spike Detector Threshold 6 x STD). Following recordings, recommended neural metrics were exported using the Axion Neural Metric software. For the recording of neuronal activity in the presence of the H current inhibitor ZD 7288, following the last recording at Day 50, half the media was removed from each well, supplemented with 50 µM ZD 7288 and re-added to each well to obtain a final concentration of 25 µM. Neuronal activity in the presence of ZD 7288 was recorded immediately for 10 minutes.

### Bulk RNA-seq

For bulk RNA-seq, neuronal progenitors were plated at a density of 170.000 cells/cm^2^ on a monolayer of primary astrocytes previously plated on GFR-Matrigel-coated plates.

At 6 weeks, samples were collected in Lysis Buffer (20 mM Tris pH 7.4, 150 mM NaCl, 5 mM MgCl_2_, 1mM DTT, 100 µg/ml Cyclohexamide, 1% TritonX-100, 25 U/ml Turbo DNAse) and total RNA was extracted from the digested lysates using Direct-zol RNA Microprep Kit (Zymo). For each sample, 3.5 µL of 50 – 250 ng/µL of RNA was fragmented in 0.5 µL SuperScript IV buffer (Invitrogen) for 4 min at 95°C. 3.5 µL of fragmented RNA, 2 µL water, 0.5 µL of 10 mM dNTPs and 0.5 µL of 5 µM of oligo-dT reverse transcription primers containing Illumina P7 sequences were mixed, and primers were annealed by heating at 65°C for 3 min then cooling to 42°C at 1°C/s. At this point, reverse transcription was carried out using SuperScript IV according to manufacturer instructions, with the addition of 0.25 µL of 40 µM of a template-switching oligo containing Illumina P5 sequences and UMIs. The reaction was incubated for 1 hour at 42°C. The product was purified using Totalpure NGS Beads and amplified using 0.5 µM i5/i7 Illumina indexed primers in Q5 High-Fidelity Master Mix (NEB). The amplified product was used for 3’ end, PE 150bp, Next Generation Sequencing with Illumina NovaSeq 6000.

### Principal Component Analysis and Volcano Plot

Spike timings for all electrodes were generated using the Axion Navigator software using default settings. Custom functions written in MATLAB were used to extract the following 20 parameters: mean spike rate, Coefficient of Variation (CoV) of Inter Spike Interval (ISI), fraction of ISIs shorter than 100ms, peak of histogram of log(ISI), electrode burst rate, electrode burst duration, spike rate within electrode bursts (first 5 spikes only), fraction of total detected spikes that are contained within an electrode burst, CoV of Inter electrode Burst Interval (IelBI), network burst rate, network burst duration, network burst participation (fraction of detected electrodes that are active during network burst), CoV of Inter network Burst Interval (InetBI), fraction of spikes that are contained within a network burst, peak frequency of Power Spectrum of electrode Burst Rate (PSelBR), peak amplitude of PSelBR, mean amplitude of PSelBR between the following 3 frequency intervals (0.1-0.35; 0.5-0.85 and 1-1.5 Hz). Electrode bursts were detected with a custom-built algorithm based on ^85^,with core parameters defined as maximum ISI = 50 ms, minimum number of spikes = 3, minimum IBI = 250 ms. Burst cores were extended in both directions if ISIs were less than 100 ms. Only bursts with at least 6 spikes were considered. Network bursts were defined by binning activity in 100 ms windows for each electrode in a well. For each bin interval, each electrode was deemed “active” if at least a spike was detected. Active status was averaged across the well for every bin and smoothed using a gaussian filter (600 ms interval, 200 ms smoothing parameter). A threshold of 40% of this smoothed activity parameter was used to detect network bursts. Only wells with at least 5 active electrodes were considered for network burst detection. Power spectrum was normalized to the integral of frequencies between 0.1 and 2 Hz. Data extracted from recordings taken during week 6 (D42, D46, D50) was averaged.

Volcano plots were created by calculating the percentage change for each parameter value in patient derived lines (DS1, DS2 and DS5) relative to the average of control lines (WT and IsoDS1). Parameters where the control average was below 0.05 were excluded from the plot. The y axis was calculated using the Mann-Whitney statistical test. Displayed data points represent average values from recordings taken during week 6 (D42, D46, D50).

### Quantification and statistical analysis

All data are presented as mean ± standard error of the mean (SEM). The nature and size of the samples, specific statistical methods carried out and exact P-values are detailed in the figure legends. Briefly, a series of normality tests (Anderson-Darling, D’Agostino & Pearson, Shapiro-Wilk, Kolmogorov-Smirnov) were performed to determine if parametric or non-parametric approaches were appropriate. For normally distributed datasets, unpaired parametric t-test was performed when comparing the effect of one variable across two datasets and ordinary One-Way ANOVA with Tukey’s multiple comparisons test was performed when comparing the effect of one variable across three or more datasets. For non-normally distributed datasets unpaired non-parametric Mann-Whitney test was performed when comparing the effect of one variable across two datasets and Kruskal-Wallis with Dunn’s multiple comparisons test was performed when comparing the effect of one variable across three or more datasets. Finally, Two-Way ANOVA with multiple comparison was performed when comparing the effect of two variables across three or more datasets.

P-values < 0.05 were considered to be statistically significant and in figures and figure legends are denoted by asterisks * (* p < 0.05, ** p < 0.01, *** p < 0.001 and **** p < 0.0001). Data presentation and statistical analyses were performed using Graphpad Prism, Version 10.5.0 (673) (Graphpad Software, San Diego, California USA, www.graphpad.com).

**Table S1.**
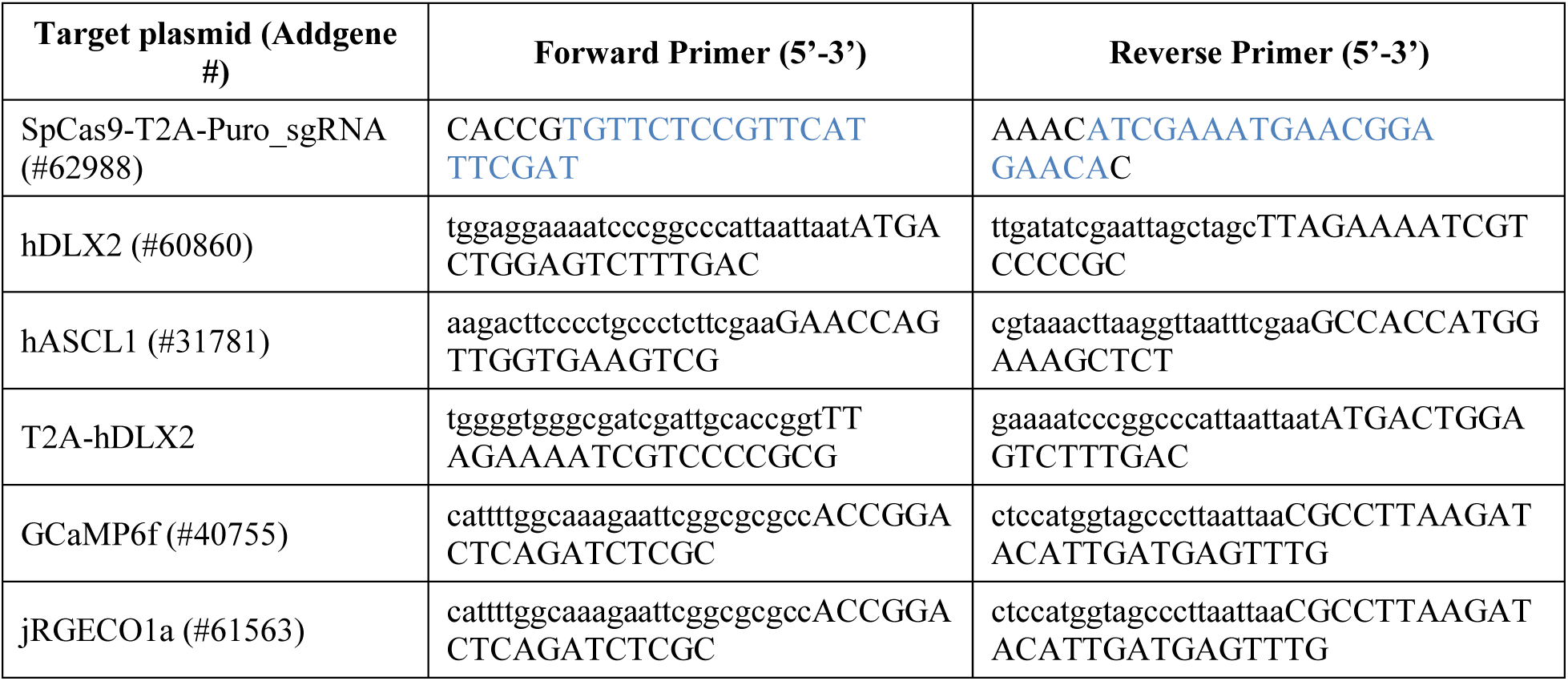
List of primers used for PCR-based cloning. Blue ink in the first set of primers highlights the 20 nucleotide sequence of the sgRNA.

**Table S2.**
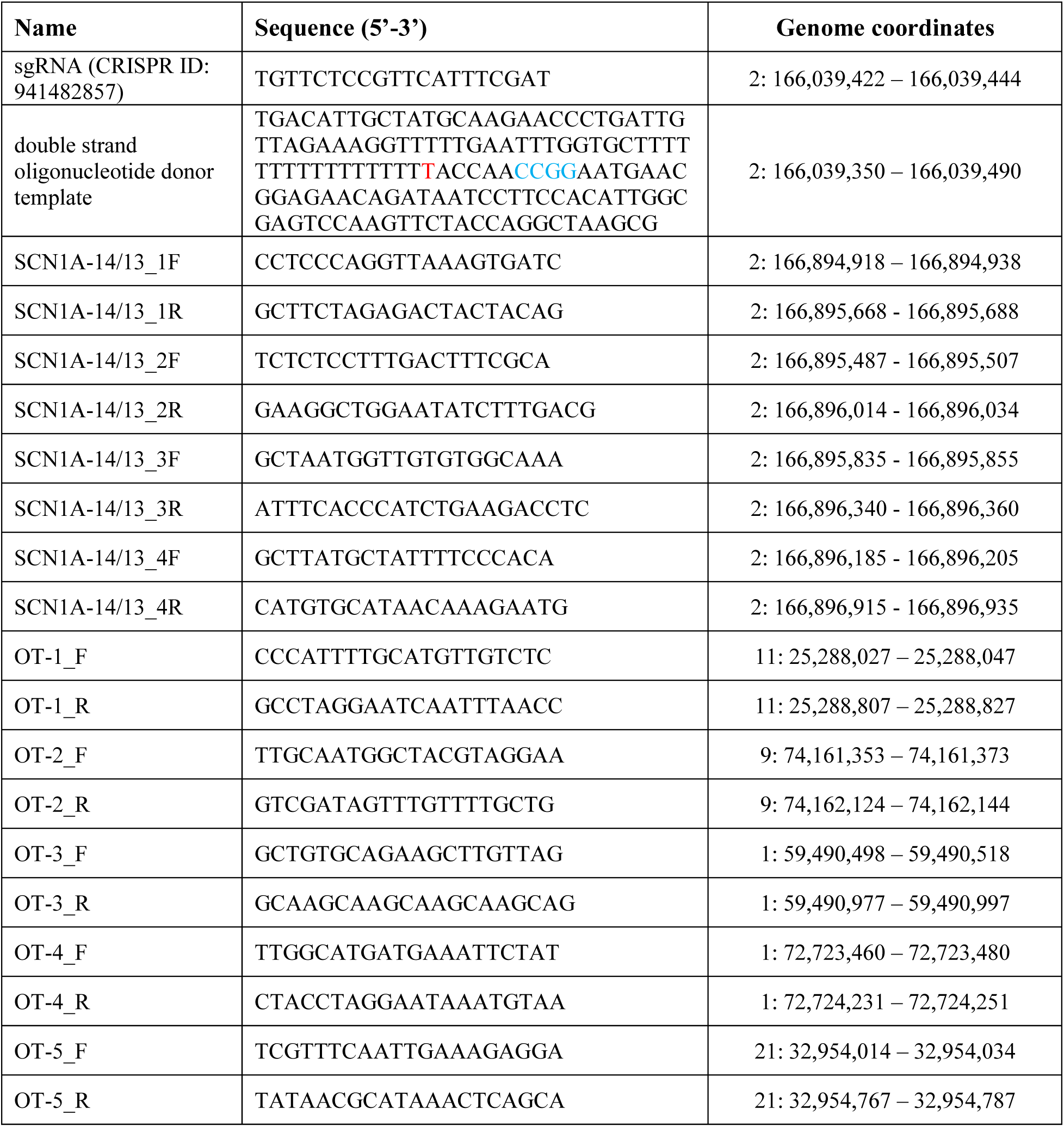
sgRNA, donor template and primer sequences used for the CRISPR/Cas9-mediated correction of the DS1 line. Red ink in the template sequence highlights the SCN1A^IVS^ ^14+3A>T^ mutation and blue ink highlights the silent MspI restriction site.

**Table S3.**
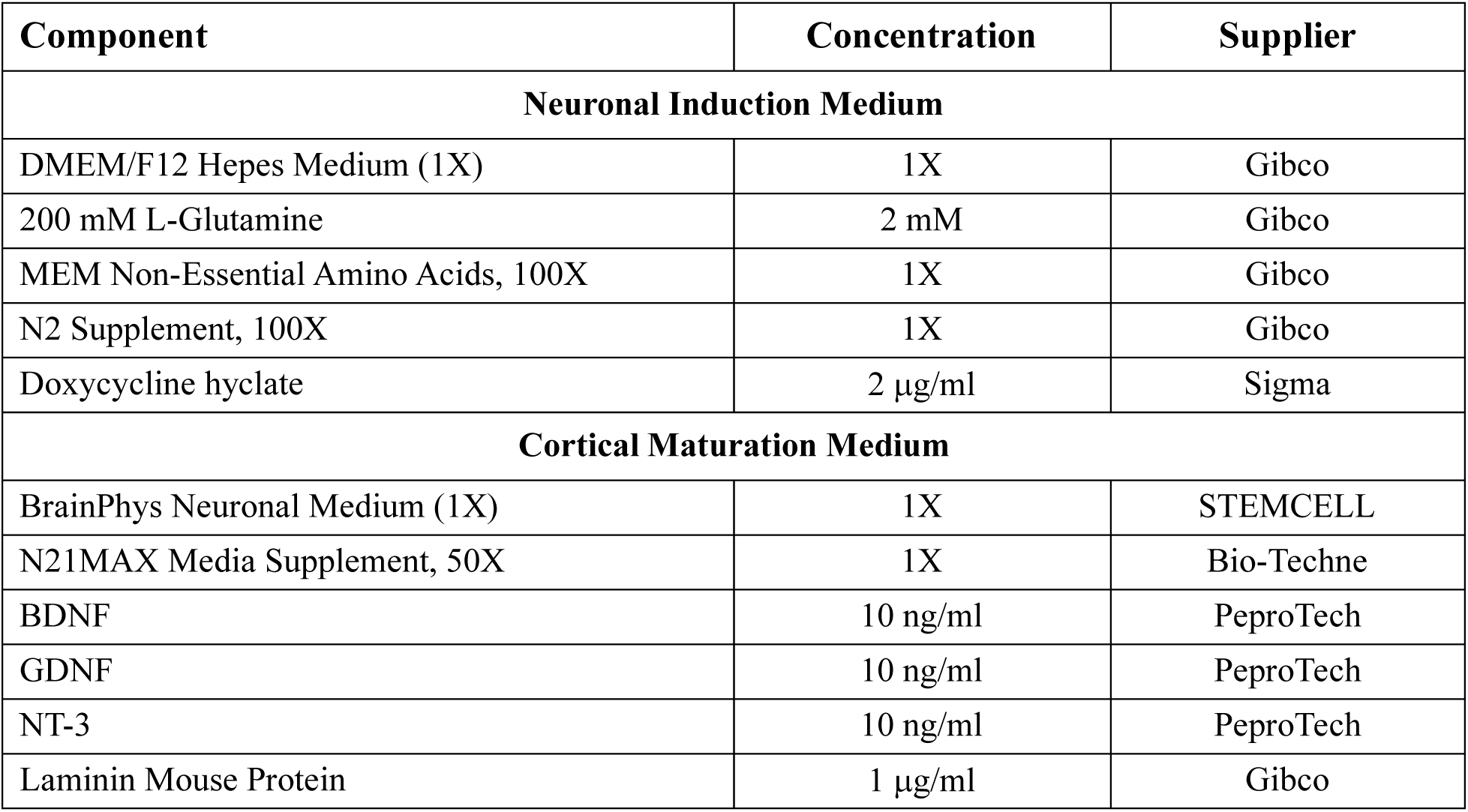
Neuronal differentiation medium compositions.

**Table S4.** List of primary and secondary antibodies used in the immunocytochemistry experiments.

| Target | Species / Fluorophore | Concentration | Supplier (Cat #) |
| --- | --- | --- | --- |
| **Primary Antibodies** |  |  |  |
| GFP | Chicken IgG | 1:500 | Abcam (ab13970) |
| Gephyrin | Mouse IgG1 | 1:200 | Synaptic Systems (147 021) |
| HCN1 | Rabbit IgG | 1:200 | Alomone (APC-056) |
| MAP2 | Rabbit IgG | 1:1000 | Antibodies-Online (ABIN1742387) |
| PSD95 | Mouse IgG2a | 1:200 | NeuroMabs (75-028) |
| RFP | Rabbit IgG | 1:500 | Abcam (ab62341) |
| TUBB3 | Mouse IgG2a | 1:1000 | R&D Systems (MAB1195) |
| **Secondary Antibodies** |  |  |  |
| Chicken | Goat – AF488 | 1:1000 | Invitrogen (A11039) |
| Mouse IgG2a | Goat – AF488 | 1:1000 | Invitrogen (A21143) |
| Mouse IgG2a | Donkey – AF647 | 1:1000 | Invitrogen (A21123) |
| Mouse IgG | Donkey – AF488 | 1:1000 | Invitrogen (A21206) |
| Rabbit IgG | Donkey – AF568 | 1:1000 | Invitrogen (A12379) |
| DNA | DAPI 405 | 1:1000 | Thermo Fisher (D1306) |

**Table S5.**
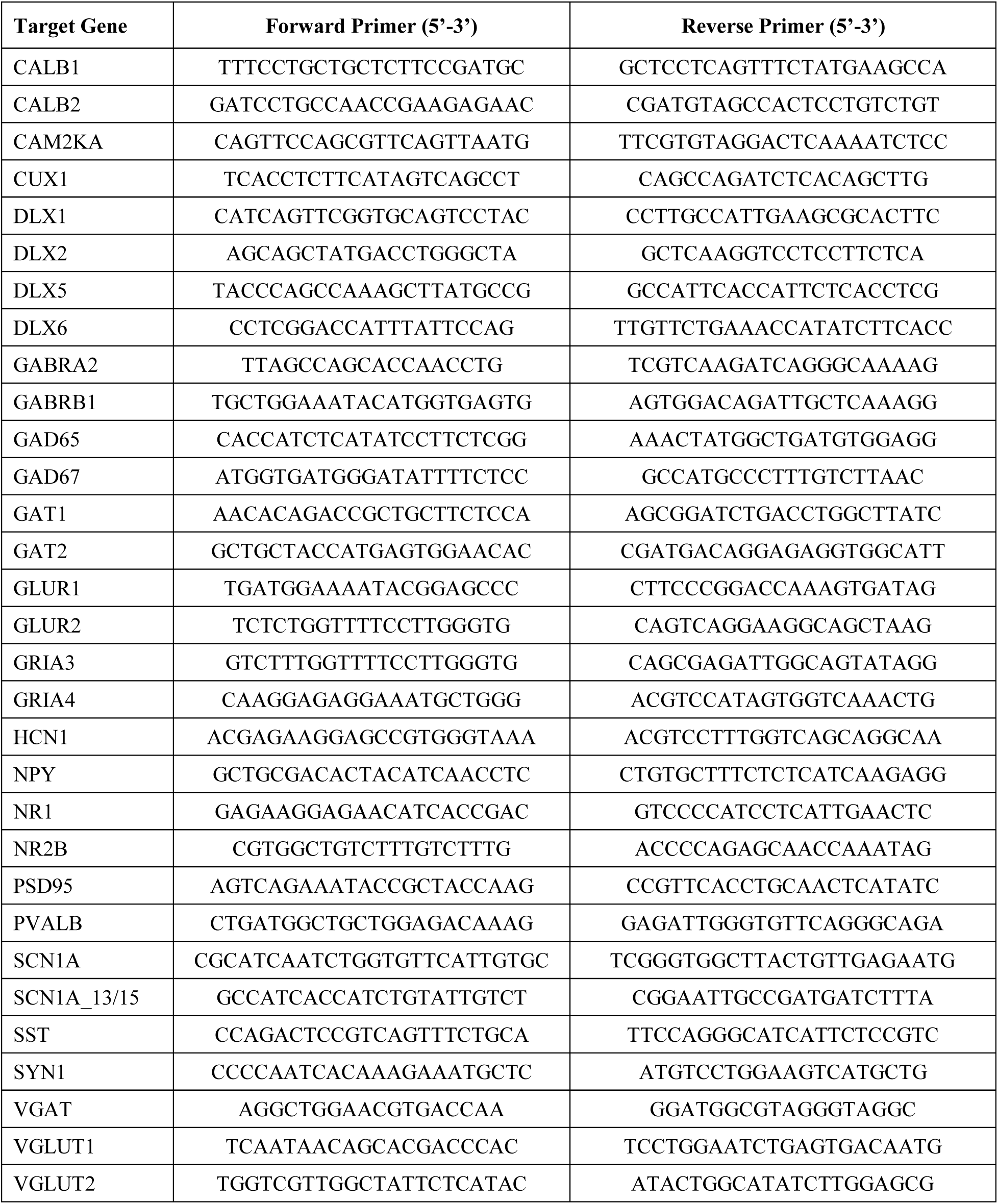
List of forward and reverse primers used for RT-qPCR analysis, in alphabetical order.

**Table S6.**
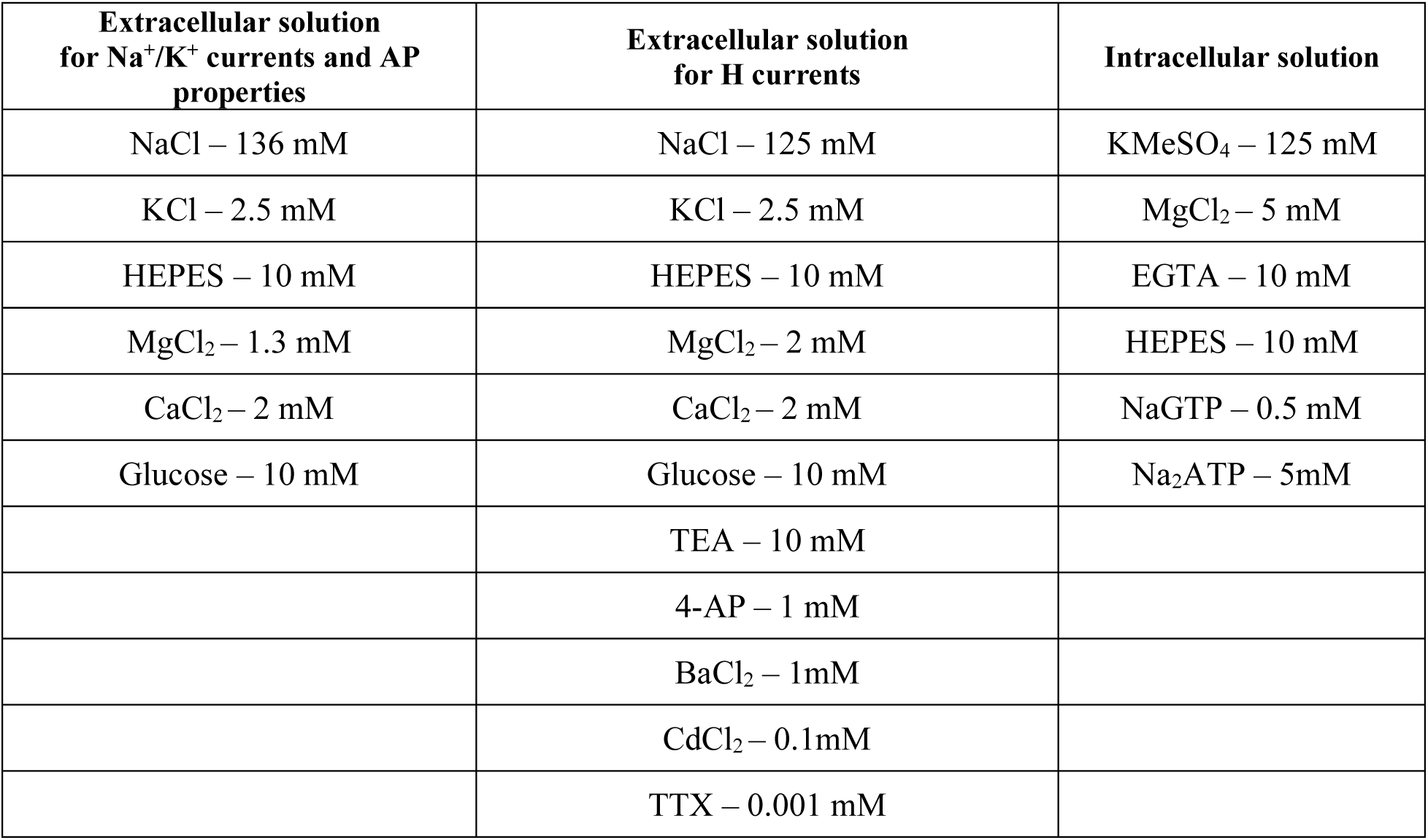
Details of intracellular/extracellular patch clamping solutions composition.

**Table S7.** Details of calcium imaging solutions composition.

| **Component** | **HEPES Buffered Saline (HBS)** | **Ca<sup>2+</sup> Imaging Buffer** |
| --- | --- | --- |
| NaCl | 136 mM | 136 mM |
| HEPES | 10 mM | 10 mM |
| Glucose | 10 mM | 10 mM |
| MgCl<sub>2</sub> | 1 mM | 1 mM |
| KCl | 2.5 mM | 8 mM |
| CaCl<sub>2</sub> | 2 mM | 4 mM |

## REFERENCES

1. C. Dravet, The core Dravet syndrome phenotype. Epilepsia 52, 3–9 (2011).

2. L. Lagae, I. Brambilla, A. Mingorance, E. Gibson, A. Battersby, Quality of life and comorbidities associated with Dravet syndrome severity: a multinational cohort survey. Dev Med Child Neurol 60, 63–72 (2018).

3. A. Brunklaus, R. Ellis, E. Reavey, G. H. Forbes, S. M. Zuberi, Prognostic, clinical and demographic features in SCN1A mutation-positive Dravet syndrome. Brain 135, 2329–2336 (2012).

4. C. Marini, I. E. Scheffer, R. Nabbout, A. Suls, P. De Jonghe, F. Zara, R. Guerrini, The genetics of Dravet syndrome. Epilepsia (2011), doi:10.1111/j.1528-1167.2011.02997.x.

5. I. Ogiwara, H. Miyamoto, N. Morita, N. Atapour, E. Mazaki, I. Inoue, T. Takeuchi, S. Itohara, Y. Yanagawa, K. Obata, T. Furuichi, T. K. Hensch, K. Yamakawa, Nav1.1 Localizes to Axons of Parvalbumin-Positive Inhibitory Interneurons: A Circuit Basis for Epileptic Seizures in Mice Carrying an Scn1a Gene Mutation. Journal of Neuroscience 27, 5903–5914 (2007).

6. F. H. Yu, M. Mantegazza, R. E. Westenbroek, C. A. Robbins, F. Kalume, K. A. Burton, W. J. Spain, G. S. McKnight, T. Scheuer, W. A. Catterall, Reduced sodium current in GABAergic interneurons in a mouse model of severe myoclonic epilepsy in infancy. Nature Neuroscience 2006 9:9 9, 1142–1149 (2006).

7. C. Tai, Y. Abe, R. E. Westenbroek, T. Scheuer, W. A. Catterall, Impaired excitability of somatostatin– and parvalbumin-expressing cortical interneurons in a mouse model of Dravet syndrome. Proc Natl Acad Sci U S A 111, E3139–E3148 (2014).

8. C. S. Cheah, F. H. Yu, R. E. Westenbroek, F. K. Kalume, J. C. Oakley, G. B. Potter, J. L. Rubenstein, W. A. Catterall, Specific deletion of NaV1.1 sodium channels in inhibitory interneurons causes seizures and premature death in a mouse model of Dravet syndrome. Proc Natl Acad Sci U S A 109, 14646–14651 (2012).

9. M. S. Martin, K. Dutt, L. A. Papale, C. M. Dubé, S. B. Dutton, G. De Haan, A. Shankar, S. Tufik, M. H. Meisler, T. Z. Baram, A. L. Goldin, A. Escayg, Altered Function of the SCN1A Voltage-gated Sodium Channel Leads to γ-Aminobutyric Acid-ergic (GABAergic) Interneuron Abnormalities. J Biol Chem 285, 9823 (2010).

10. A. M. Mistry, C. H. Thompson, A. R. Miller, C. G. Vanoye, A. L. George, J. A. Kearney, Strain– and Age-dependent Hippocampal Neuron Sodium Currents Correlate with Epilepsy Severity in Dravet Syndrome Mice. Neurobiol Dis 65, 1 (2014).

11. Y. Almog, S. Fadila, M. Brusel, A. Mavashov, K. Anderson, M. Rubinstein, Developmental alterations in firing properties of hippocampal CA1 inhibitory and excitatory neurons in a mouse model of Dravet syndrome. Neurobiol Dis 148, 105209 (2021).

12. Y. Liu, L. F. Lopez-Santiago, Y. Yuan, J. M. Jones, H. Zhang, H. A. O’Malley, G. A. Patino, J. E. O’Brien, R. Rusconi, A. Gupta, R. C. Thompson, M. R. Natowicz, M. H. Meisler, L. L. Isom, J. M. Parent, Dravet syndrome patient-derived neurons suggest a novel epilepsy mechanism. Ann Neurol (2013), doi:10.1002/ana.23897.

13. J. Jiao, Y. Yang, Y. Shi, J. Chen, R. Gao, Y. Fan, H. Yao, W. Liao, X. F. Sun, S. Gao, Modeling Dravet syndrome using induced pluripotent stem cells (iPSCs) and directly converted neurons. Hum Mol Genet 22, 4241–4252 (2013).

14. E. J. H. Van Hugte, E. I. Lewerissa, K. M. Wu, N. Scheefhals, G. Parodi, T. W. Van Voorst, S. Puvogel, N. Kogo, J. M. Keller, M. Frega, D. Schubert, H. J. Schelhaas, J. Verhoeven, M. Majoie, H. Van Bokhoven, N. N. Kasri, SCN1A-deficient excitatory neuronal networks display mutation-specific phenotypes. Brain 146, 5153 (2023).

15. M. Biel, C. Wahl-Schott, S. Michalakis, X. Zong, Hyperpolarization-activated cation channels: From genes to function. Physiol Rev 89, 847–885 (2009).

16. E. E. Benarroch, HCN channels: Function and clinical implications. Neurology 80, 304–310 (2013).

17. C. R. Frasier, H. Zhang, J. Offord, L. T. Dang, D. S. Auerbach, H. Shi, C. Chen, A. M. Goldman, L. L. Eckhardt, V. J. Bezzerides, J. M. Parent, L. L. Isom, Channelopathy as a SUDEP Biomarker in Dravet Syndrome Patient-Derived Cardiac Myocytes. Stem Cell Reports 11, 626 (2018).

18. M. S. Fernandopulle, R. Prestil, C. Grunseich, C. Wang, L. Gan, M. E. Ward, Transcription Factor–Mediated Differentiation of Human iPSCs into Neurons. Curr Protoc Cell Biol (2018), doi:10.1002/cpcb.51.

19. N. Yang, S. Chanda, S. Marro, Y. H. Ng, J. A. Janas, D. Haag, C. E. Ang, Y. Tang, Q. Flores, M. Mall, O. Wapinski, M. Li, H. Ahlenius, J. L. Rubenstein, H. Y. Chang, A. A. Buylla, T. C. Südhof, M. Wernig, Generation of pure GABAergic neurons by transcription factor programming. Nat Methods (2017), doi:10.1038/nmeth.4291.

20. T. E. Bakken, N. L. Jorstad, Q. Hu, B. B. Lake, W. Tian, B. E. Kalmbach, M. Crow, R. D. Hodge, F. M. Krienen, S. A. Sorensen, J. Eggermont, Z. Yao, B. D. Aevermann, A. I. Aldridge, A. Bartlett, D. Bertagnolli, T. Casper, R. G. Castanon, K. Crichton, T. L. Daigle, R. Dalley, N. Dee, N. Dembrow, D. Diep, S. L. Ding, W. Dong, R. Fang, S. Fischer, M. Goldman, J. Goldy, L. T. Graybuck, B. R. Herb, X. Hou, J. Kancherla, M. Kroll, K. Lathia, B. van Lew, Y. E. Li, C. S. Liu, H. Liu, J. D. Lucero, A. Mahurkar, D. McMillen, J. A. Miller, M. Moussa, J. R. Nery, P. R. Nicovich, S. Y. Niu, J. Orvis, J. K. Osteen, S. Owen, C. R. Palmer, T. Pham, N. Plongthongkum, O. Poirion, N. M. Reed, C. Rimorin, A. Rivkin, W. J. Romanow, A. E. Sedeño-Cortés, K. Siletti, S. Somasundaram, J. Sulc, M. Tieu, A. Torkelson, H. Tung, X. Wang, F. Xie, A. M. Yanny, R. Zhang, S. A. Ament, M. M. Behrens, H. C. Bravo, J. Chun, A. Dobin, J. Gillis, R. Hertzano, P. R. Hof, T. Höllt, G. D. Horwitz, C. D. Keene, P. V. Kharchenko, A. L. Ko, B. P. Lelieveldt, C. Luo, E. A. Mukamel, A. Pinto-Duarte, S. Preissl, A. Regev, B. Ren, R. H. Scheuermann, K. Smith, W. J. Spain, O. R. White, C. Koch, M. Hawrylycz, B. Tasic, E. Z. Macosko, S. A. McCarroll, J. T. Ting, H. Zeng, K. Zhang, G. Feng, J. R. Ecker, S. Linnarsson, E. S. Lein, Comparative cellular analysis of motor cortex in human, marmoset and mouse. Nature 2021 598:7879 598, 111–119 (2021).

20. D. Džaja, A. Hladnik, I. Bičanić, M. Baković, Z. Petanjek, Neocortical calretinin neurons in primates: increase in proportion and microcircuitry structure. Front Neuroanat 8, 103 (2014).

21. Z. Friedenberger, E. Harkin, K. Tóth, R. Naud, Silences, spikes and bursts: Three-part knot of the neural code. J Physiol 601, 5165–5193 (2023).

22. D. A. Eisner, E. Neher, H. Taschenberger, G. Smith, Physiology of intracellular calcium buffering. Physiol Rev 103, 2767 (2023).

23. I. E. Scheffer, R. Nabbout, SCN1A-related phenotypes: Epilepsy and beyond. Epilepsia 60, S17–S24 (2019).

24. I. M. de Lange, M. J. Koudijs, R. van ‘t Slot, B. Gunning, A. C. M. Sonsma, L. J. J. M. van Gemert, F. Mulder, E. C. Carbo, M. J. A. van Kempen, N. E. Verbeek, I. J. Nijman, R. F. Ernst, S. M. C. Savelberg, N. V. A. M. Knoers, E. H. Brilstra, B. P. C. Koeleman, Mosaicism of de novo pathogenic SCN1A variants in epilepsy is a frequent phenomenon that correlates with variable phenotypes. Epilepsia 59, 690–703 (2018).

25. R. Guerrini, E. Cellini, D. Mei, T. Metitieri, C. Petrelli, D. Pucatti, C. Marini, N. Zamponi, Variable epilepsy phenotypes associated with a familial intragenic deletion of the SCN1A gene. Epilepsia 51, 2474–2477 (2010).

26. H. Martins Custodio, L. M. Clayton, R. Bellampalli, S. Pagni, K. Silvennoinen, R. Caswell, J. C. Ambrose, P. Arumugam, R. Bevers, M. Bleda, F. Boardman-Pretty, C. R. Boustred, H. Brittain, M. A. Brown, M. J. Caulfield, G. C. Chan, A. Giess, J. N. Griffin, A. Hamblin, S. Henderson, T. J. P. Hubbard, R. Jackson, L. J. Jones, D. Kasperaviciute, M. Kayikci, A. Kousathanas, L. Lahnstein, A. Lakey, S. E. A. Leigh, I. U. S. Leong, J. F. Lopez, F. Maleady-Crowe, M. Mcentagart, F. Minneci, J. Mitchell, L. Moutsianas, M. Mueller, N. Murugaesu, A. C. Need, P. O’donovan, C. A. Odhams, C. Patch, D. Perez-Gil, M. B. Pereira, J. Pullinger, T. Rahim, A. Rendon, T. Rogers, K. Savage, K. Sawant, R. H. Scott, A. Siddiq, A. Sieghart, S. C. Smith, A. Sosinsky, A. Stuckey, M. Tanguy, A. L. T. Tavares, E. R. A. Thomas, S. R. Thompson, A. Tucci, M. J. Welland, E. Williams, K. Witkowska, S. M. Wood, M. Zarowiecki, A. Brunklaus, R. Guerrini, B. P. C. Koeleman, J. R. Lemke, R. S. Møller, I. E. Scheffer, S. Weckhuysen, F. Zara, S. Zuberi, K. Kuchenbaecker, S. Balestrini, J. D. Mills, S. M. Sisodiya, Widespread genomic influences on phenotype in Dravet syndrome, a ‘monogenic’ condition. Brain 146, 3885 (2023).

27. I. M. de Lange, W. Weuring, R. van ‘t Slot, B. Gunning, A. C. M. Sonsma, M. McCormack, C. de Kovel, L. J. J. M. van Gemert, F. Mulder, M. J. A. van Kempen, N. V. A. M. Knoers, E. H. Brilstra, B. P. C. Koeleman, Influence of common SCN1A promoter variants on the severity of SCN1A-related phenotypes. Mol Genet Genomic Med 7, e00727 (2019).

28. N. Higurashi, T. Uchida, C. Lossin, Y. Misumi, Y. Okada, W. Akamatsu, Y. Imaizumi, B. Zhang, K. Nabeshima, M. X. Mori, S. Katsurabayashi, Y. Shirasaka, H. Okano, S. Hirose, A human Dravet syndrome model from patient induced pluripotent stem cells. Mol Brain 6, 19 (2013).

29. H. W. Kim, Z. Quan, Y. B. Kim, E. Cheong, H. D. Kim, M. Cho, J. Jang, Y. R. Yoo, J. S. Lee, J. H. Kim, Y. I. Kim, D. S. Kim, H. C. Kang, Differential effects on sodium current impairments by distinct SCN1A mutations in GABAergic neurons derived from Dravet syndrome patients. Brain Dev 40, 287–298 (2018).

30. P. W. Marshall, W. Rouse, I. Briggs, R. B. Hargreaves, S. D. Mills, B. J. McLoughlin, ICI D7288, a novel sinoatrial node modulator. J Cardiovasc Pharmacol 21, 902–906 (1993).

31. S. Gasparini, D. DiFrancesco, Action of the hyperpolarization-activated current (I(h)) blocker ZD 7288 in hippocampal CA1 neurons. Pflugers Arch 435, 99–106 (1997).

32. T. O. Satoh, M. Yamada, A bradycardiac agent ZD7288 blocks the hyperpolarization-activated current (Ih) in retinal rod photoreceptors. Neuropharmacology 39, 1284–1291 (2000).

33. Y. Inaba, G. Biagini, M. Avoli, The H current blocker ZD7288 decreases epileptiform hyperexcitability in the rat neocortex by depressing synaptic transmission. Neuropharmacology 51, 681–691 (2006).

34. X. Wu, L. Liao, X. Liu, F. Luo, T. Yang, C. Li, Is ZD7288 a selective blocker of hyperpolarization-activated cyclic nucleotide-gated channel currents? Channels 6, 438 (2012).

35. J. Liu, C. Gao, W. Chen, W. Ma, X. Li, Y. Shi, H. Zhang, L. Zhang, Y. Long, H. Xu, X. Guo, S. Deng, X. Yan, D. Yu, G. Pan, Y. Chen, L. Lai, W. Liao, Z. Li, CRISPR/Cas9 facilitates investigation of neural circuit disease using human iPSCs: mechanism of epilepsy caused by an SCN1A loss-of-function mutation. Transl Psychiatry 6, e703 (2016).

36. A. M. De Stasi, P. Farisello, I. Marcon, S. Cavallari, A. Forli, D. Vecchia, G. Losi, M. Mantegazza, S. Panzeri, G. Carmignoto, A. Bacci, T. Fellin, Unaltered Network Activity and Interneuronal Firing During Spontaneous Cortical Dynamics In Vivo in a Mouse Model of Severe Myoclonic Epilepsy of Infancy. Cerebral Cortex (New York, NY) 26, 1778 (2016).

37. M. Favero, N. P. Sotuyo, E. Lopez, J. A. Kearney, E. M. Goldberg, A Transient Developmental Window of Fast-Spiking Interneuron Dysfunction in a Mouse Model of Dravet Syndrome. Journal of Neuroscience 38, 7912–7927 (2018).

38. C. H. Tran, M. Vaiana, J. Nakuci, A. Somarowthu, K. M. Goff, N. Goldstein, P. Murthy, S. F. Muldoon, E. M. Goldberg, Interneuron Desynchronization Precedes Seizures in a Mouse Model of Dravet Syndrome. Journal of Neuroscience 40, 2764–2775 (2020).

39. Y. Sun, S. P. Paşca, T. Portmann, C. Goold, K. A. Worringer, W. Guan, K. C. Chan, H. Gai, D. Vogt, Y. J. J. Chen, R. Mao, K. Chan, J. L. R. Rubenstein, D. V. Madison, J. Hallmayer, W. M. Froehlich-Santino, J. A. Bernstein, R. E. Dolmetsch, A deleterious Nav1.1 mutation selectively impairs telencephalic inhibitory neurons derived from Dravet Syndrome patients. Elife (2016), doi:10.7554/eLife.13073.

40. T. Arimitsu, M. Nuriya, K. Ikeda, T. Takahashi, M. Yasui, Activity-dependent regulation of HCN1 protein in cortical neurons. Biochem Biophys Res Commun 387, 87–91 (2009).

41. S. Jung, T. D. Jones, J. N. Lugo, A. H. Sheerin, J. W. Miller, R. D’Ambrosio, A. E. Anderson, N. P. Poolos, Progressive Dendritic HCN Channelopathy during Epileptogenesis in the Rat Pilocarpine Model of Epilepsy. The Journal of Neuroscience 27, 13012 (2007).

42. B. E. L. Adams, C. A. Reid, D. Myers, C. Ng, K. Powell, A. M. Phillips, T. Zheng, T. J. O’Brien, D. A. Williams, Excitotoxic-mediated transcriptional decreases in HCN2 channel function increase network excitability in CA1. Exp Neurol 219, 249–257 (2009).

43. S. Jung, L. N. Warner, J. Pitsch, A. J. Becker, N. P. Poolos, Rapid Loss of Dendritic HCN Channel Expression in Hippocampal Pyramidal Neurons following Status Epilepticus. The Journal of Neuroscience 31, 14291 (2011).

44. K. Chen, I. Aradi, N. Thon, M. Eghbal-Ahmadi, T. Z. Baram, I. Soltesz, Persistently modified h-channels after complex febrile seizures convert the seizure-induced enhancement of inhibition to hyperexcitability. Nat Med 7, 331 (2001).

45. J. Dyhrfjeld-Johnsen, R. J. Morgan, C. Földy, I. Soltesz, Upregulated H-Current in Hyperexcitable CA1 Dendrites after Febrile Seizures. Front Cell Neurosci 2, 2 (2008).

46. M. M. Shah, A. E. Anderson, V. Leung, X. Lin, D. Johnston, Seizure-Induced Plasticity of h Channels in Entorhinal Cortical Layer III Pyramidal Neurons. Neuron 44, 495 (2004).

47. K. Zhang, B. W. Peng, R. M. Sanchez, Decreased IH in hippocampal area CA1 pyramidal neurons after perinatal seizure-inducing hypoxia. Epilepsia 47, 1023–1028 (2006).

48. U. Strauss, M. H. P. Kole, A. U. Bräuer, J. Pahnke, R. Bajorat, A. Rolfs, R. Nitsch, R. A. Deisz, An impaired neocortical Ih is associated with enhanced excitability and absence epilepsy. Eur J Neurosci 19, 3048–3058 (2004).

49. M. H. P. Kole, A. U. Bräuer, G. J. Stuart, Inherited cortical HCN1 channel loss amplifies dendritic calcium electrogenesis and burst firing in a rat absence epilepsy model. J Physiol 578, 507 (2006).

50. C. Nava, C. Dalle, A. Rastetter, P. Striano, C. G. F. De Kovel, R. Nabbout, C. Cancès, D. Ville, E. H. Brilstra, G. Gobbi, E. Raffo, D. Bouteiller, Y. Marie, O. Trouillard, A. Robbiano, B. Keren, D. Agher, E. Roze, S. Lesage, A. Nicolas, A. Brice, M. Baulac, C. Vogt, N. El Hajj, E. Schneider, A. Suls, S. Weckhuysen, P. Gormley, A. E. Lehesjoki, P. De Jonghe, I. Helbig, S. Baulac, F. Zara, B. P. C. Koeleman, T. Haaf, E. Leguern, C. Depienne, De novo mutations in HCN1 cause early infantile epileptic encephalopathy. Nature Genetics 2014 46:6 46, 640–645 (2014).

51. C. Marini, A. Porro, A. Rastetter, C. Dalle, I. Rivolta, D. Bauer, R. Oegema, C. Nava, E. Parrini, D. Mei, C. Mercer, R. Dhamija, C. Chambers, C. Coubes, J. Thévenon, P. Kuentz, S. Julia, L. Pasquier, C. Dubourg, W. Carré, A. Rosati, F. Melani, T. Pisano, M. Giardino, A. M. Innes, Y. Alembik, S. Scheidecker, M. Santos, S. Figueiroa, C. Garrido, C. Fusco, D. Frattini, C. Spagnoli, A. Binda, T. Granata, F. Ragona, E. Freri, S. Franceschetti, L. Canafoglia, B. Castellotti, C. Gellera, R. Milanesi, M. M. Mancardi, D. R. Clark, F. Kok, K. L. Helbig, S. Ichikawa, L. Sadler, J. Neupauerová, P. Laššuthova, K. Štěrbová, A. Laridon, E. Brilstra, B. Koeleman, J. R. Lemke, F. Zara, P. Striano, J. Soblet, G. Smits, N. Deconinck, A. Barbuti, D. Difrancesco, E. Leguern, R. Guerrini, B. Santoro, K. Hamacher, G. Thiel, A. Moroni, J. C. Difrancesco, C. Depienne, HCN1 mutation spectrum: from neonatal epileptic encephalopathy to benign generalized epilepsy and beyond. Brain 141, 3160–3178 (2018).

52. D. Steel, J. D. Symonds, S. M. Zuberi, A. Brunklaus, Dravet syndrome and its mimics: Beyond SCN1A. Epilepsia 58, 1807–1816 (2017).

53. R. Guerrini, A. Belmonte, P. Genton, Antiepileptic Drug-Induced Worsening of Seizures in Children. Epilepsia 39, S2–S10 (1998).

54. N. A. Hawkins, L. L. Anderson, T. S. Gertler, L. Laux, A. L. George, J. A. Kearney, Screening of conventional anticonvulsants in a genetic mouse model of epilepsy. Ann Clin Transl Neurol 4, 326–339 (2017).

55. J. Lehnhoff, U. Strauss, S. Wierschke, S. Grosser, E. Pollali, U. C. Schneider, M. Holtkamp, C. Dehnicke, R. A. Deisz, The anticonvulsant lamotrigine enhances Ih in layer 2/3 neocortical pyramidal neurons of patients with pharmacoresistant epilepsy. Neuropharmacology 144, 58–69 (2019).

56. L. E. Bleakley, C. E. McKenzie, C. A. Reid, Efficacy of antiseizure medication in a mouse model of HCN1 developmental and epileptic encephalopathy. Epilepsia 64, e1 (2022).

57. N. P. Poolos, M. Migliore, D. Johnston, Pharmacological upregulation of h-channels reduces the excitability of pyramidal neuron dendrites. Nature Neuroscience 2002 5:8 5, 767–774 (2002).

58. A. Omrani, T. Van Der Vaart, E. Mientjes, G. M. Van Woerden, M. R. Hojjati, K. W. Li, D. H. Gutmann, C. N. Levelt, A. B. Smit, A. J. Silva, S. A. Kushner, Y. Elgersma, HCN channels are a novel therapeutic target for cognitive dysfunction in Neurofibromatosis type 1. Mol Psychiatry 20, 1311 (2015).

59. T. Berger, H. R. Lüscher, Associative somatodendritic interaction in layer V pyramidal neurons is not affected by the antiepileptic drug lamotrigine. European Journal of Neuroscience 20, 1688–1693 (2004).

60. M. Rubinstein, R. E. Westenbroek, F. H. Yu, C. J. Jones, T. Scheuer, W. A. Catterall, Genetic Background Modulates Impaired Excitability of Inhibitory Neurons in a Mouse Model of Dravet Syndrome. Neurobiol Dis 73, 106 (2015).

61. A. Lörincz, T. Notomi, G. Tamás, R. Shigemoto, Z. Nusser, Polarized and compartment-dependent distribution of HCN1 in pyramidal cell dendrites. Nature Neuroscience 2002 5:11 5, 1185–1193 (2002).

62. B. Santoro, S. Chen, A. Lüthi, P. Pavlidis, G. P. Shumyatsky, G. R. Tibbs, S. A. Siegelbaum, Molecular and Functional Heterogeneity of Hyperpolarization-Activated Pacemaker Channels in the Mouse CNS. The Journal of Neuroscience 20, 5264 (2000).

63. H. Zeng, E. H. Shen, J. G. Hohmann, S. W. Oh, A. Bernard, J. J. Royall, K. J. Glattfelder, S. M. Sunkin, J. A. Morris, A. L. Guillozet-Bongaarts, K. A. Smith, A. J. Ebbert, B. Swanson, L. Kuan, D. T. Page, C. C. Overly, E. S. Lein, M. J. Hawrylycz, P. R. Hof, T. M. Hyde, J. E. Kleinman, A. R. Jones, Large-scale cellular-resolution gene profiling in human neocortex reveals species-specific molecular signatures. Cell 149, 483 (2012).

64. M. E. Larkum, J. Waters, B. Sakmann, F. Helmchen, Dendritic Spikes in Apical Dendrites of Neocortical Layer 2/3 Pyramidal Neurons. The Journal of Neuroscience 27, 8999 (2007).

65. B. N. Routh, R. K. Rathour, M. E. Baumgardner, B. E. Kalmbach, D. Johnston, D. H. Brager, Increased transient Na+ conductance and action potential output in layer 2/3 prefrontal cortex neurons of the fmr1−/y mouse. J Physiol 595, 4431 (2017).

66. K. I. Van Aerde, D. Feldmeyer, Morphological and Physiological Characterization of Pyramidal Neuron Subtypes in Rat Medial Prefrontal Cortex. Cerebral Cortex 25, 788–805 (2015).

67. B. E. Kalmbach, A. Buchin, B. Long, J. Close, A. Nandi, J. A. Miller, T. E. Bakken, R. D. Hodge, P. Chong, R. de Frates, K. Dai, Z. Maltzer, P. R. Nicovich, C. D. Keene, D. L. Silbergeld, R. P. Gwinn, C. Cobbs, A. L. Ko, J. G. Ojemann, C. Koch, C. A. Anastassiou, E. S. Lein, J. T. Ting, h-Channels Contribute to Divergent Intrinsic Membrane Properties of Supragranular Pyramidal Neurons in Human versus Mouse Cerebral Cortex. Neuron 100, 1194–1208.e5 (2018).

68. S. P. Jones, N. O’Neill, J. C. Carpenter, S. Muggeo, G. Colasante, D. M. Kullmann, G. Lignani, Early developmental alterations of CA1 pyramidal cells in Dravet syndrome. Neurobiol Dis 201, 106688 (2024).

69. K. Hennis, M. Biel, C. Wahl-Schott, S. Fenske, Beyond pacemaking: HCN channels in sinoatrial node function. Prog Biophys Mol Biol 166, 51–60 (2021).

70. N. Li, T. A. Csepe, B. J. Hansen, H. Dobrzynski, R. S. D. Higgins, A. Kilic, P. J. Mohler, P. M. L. Janssen, M. R. Rosen, B. J. Biesiadecki, V. V. Fedorov, Molecular Mapping of Sinoatrial Node HCN Channel Expression in the Human Heart. Circ Arrhythm Electrophysiol 8, 1219–1227 (2015).

71. O. Devinsky, D. C. Hesdorffer, D. J. Thurman, S. Lhatoo, G. Richerson, Sudden unexpected death in epilepsy: epidemiology, mechanisms, and prevention. Lancet Neurol 15, 1075–1088 (2016).

72. S. Y. Lyu, S. O. Nam, Y. J. Lee, G. Kim, Y. A. Kim, J. Kong, A. Ko, Y. M. Kim, G. M. Yeon, Longitudinal change of cardiac electrical and autonomic function and potential risk factors in children with dravet syndrome. Epilepsy Res 152, 11–17 (2019).

73. S. Shmuely, R. Surges, R. M. Helling, W. B. Gunning, E. H. Brilstra, J. S. Verhoeven, J. H. Cross, S. M. Sisodiya, H. L. Tan, J. W. Sander, R. D. Thijs, Cardiac arrhythmias in Dravet syndrome: an observational multicenter study. Ann Clin Transl Neurol 7, 462–473 (2020).

74. N. Villas, M. A. Meskis, S. Goodliffe, Dravet syndrome: Characteristics, comorbidities, and caregiver concerns. Epilepsy & Behavior 74, 81–86 (2017).

75. M. Coll, C. Allegue, S. Partemi, J. Mates, B. Del Olmo, O. Campuzano, V. Pascali, A. Iglesias, P. Striano, A. Oliva, R. Brugada, Genetic investigation of sudden unexpected death in epilepsy cohort by panel target resequencing. Int J Legal Med 130, 331–339 (2016).

76. E. Tu, L. Waterhouse, J. Duflou, R. D. Bagnall, C. Semsarian, Genetic Analysis of Hyperpolarization-Activated Cyclic Nucleotide-Gated Cation Channels in Sudden Unexpected Death in Epilepsy Cases. Brain Pathology 21, 692 (2011).

77. F. Hofmann, L. Fabritz, J. Stieber, J. Schmitt, P. Kirchhof, A. Ludwig, S. Herrmann, Ventricular HCN channels decrease the repolarization reserve in the hypertrophic heart. Cardiovasc Res 95, 317–326 (2012).

78. Y. Kuwabara, K. Kuwahara, M. Takano, H. Kinoshita, Y. Arai, S. Yasuno, Y. Nakagawa, S. Igata, S. Usami, T. Minami, Y. Yamada, K. Nakao, C. Yamada, J. Shibata, T. Nishikimi, K. Ueshima, K. Nakao, Increased expression of HCN channels in the ventricular myocardium contributes to enhanced arrhythmicity in mouse failing hearts. J Am Heart Assoc 2 (2013), doi:10.1161/JAHA.113.000150/ASSET/B437C882-C179-470A-B3D6-3A77DDE73DDE/ASSETS/IMAGES/LARGE/JAH3227-FIG-0009.JPG.

79. P. Dias, C. M. Terracciano, Hyperpolarization-Activated Cyclic Nucleotide-Gated Channels and Ventricular Arrhythmias in Heart Failure: A Novel Target for Therapy? J Am Heart Assoc 2, 287 (2013).

80. X. Li, N. Lobo, C. A. Bauser, M. J. Fraser, The minimum internal and external sequence requirements for transposition of the eukaryotic transformation vector piggyBac. Molecular Genetics and Genomics 266, 190–198 (2001).

81. C. Wang, M. E. Ward, R. Chen, K. Liu, T. E. Tracy, X. Chen, M. Xie, P. D. Sohn, C. Ludwig, A. Meyer-Franke, C. M. Karch, S. Ding, L. Gan, Scalable Production of iPSC-Derived Human Neurons to Identify Tau-Lowering Compounds by High-Content Screening. Stem Cell Reports (2017), doi:10.1016/j.stemcr.2017.08.019.

82. K. J. Livak, T. D. Schmittgen, Analysis of Relative Gene Expression Data Using Real-Time Quantitative PCR and the 2−ΔΔCT Method. Methods 25, 402–408 (2001).

83. Z. Sun, T. C. Südhof, A simple Ca2+-imaging approach to neural network analyses in cultured neurons. J Neurosci Methods 349 (2021), doi:10.1016/J.JNEUMETH.2020.109041.

84. L. L. Bologna, V. Pasquale, M. Garofalo, M. Gandolfo, P. L. Baljon, A. Maccione, S. Martinoia, M. Chiappalone, Investigating neuronal activity by SPYCODE multi-channel data analyzer. Neural Networks 23, 685–697 (2010).

